# Data-Driven 3D Shape Analysis Reveals Cell Shape-Fate Relationships in Zebrafish Lateral Line Neuromasts

**DOI:** 10.1101/2023.08.09.552694

**Authors:** Madeleine N. Hewitt, Iván A. Cruz, David W. Raible

## Abstract

Cell shape is a powerful readout of cell state, fate, and function. With the advent of sophisticated microscopes, image segmentation algorithms, and numerical shape representations, it is becoming more feasible to study cell shape in developing tissues. However, few studies have analyzed cell shape in three dimensions in living, intact organisms. Here, we took advantage of the favorable imaging qualities of zebrafish lateral line neuromasts to generate a dataset of high resolution images with labeled cells and nuclei. Using a custom Python-based workflow, we performed semi-automated, 3D cell and nucleus segmentation. We then used spherical harmonics and principal components analysis to distill neuromast cell and nuclear shape variation into several interpretable, biologically meaningful parameters. We found that neuromast cell and nuclear shapes vary with cell location and identity. The distinction between hair cells and support cells was discrete and accounted for much of the variation in neuromast cell and nucleus shape, which allowed us to train classifiers to predict hair cell identity from cell and nucleus shape features. Using markers for support cell subpopulations, we found that support cell subtypes also had different shapes from each other; however, shape features did not distinguish as sharply between support cell subtypes, suggesting that support cells vary continuously in shape. To investigate the effects of genetic perturbation that results in loss of a cell type on neuromast cell shape, we examined *atoh1a* mutants that lack hair cells. We found that neuromasts from *atoh1a* mutants lacked the cell shape phenotype associated with hair cells, but did not exhibit a mutant-specific cell shape. Our results demonstrate the utility of using 3D cell shape features to characterize, compare, and classify cells in a living, developing organism.

## Introduction

The shape of a cell emerges from a complex interplay between fate, function, and external forces.^1^ Quantifying cell shape during development has provided mechanistic insight into morphogenetic processes, such as germ-band extension in *Drosophila.*^2,3^ Modifying cell shape in culture can influence cell division and death,^4^ as well as differentiation.^5,6^ However, most studies of cell shape *in vivo* have used descriptive features (such as volume and sphericity), which may not capture subtle or multifactorial variations in shape between cell types, especially in tissues with complex, 3D organization. More complete 2D shape representations have been used to study cell motility and migration in culture,^7,8^ but these methods cannot be easily generalized to 3D shapes. To fully appreciate the diversity of cell shape states during development and understand the relationship between cell shape and fate, robust 3D cell shape analysis methods must be applied to living, developing organisms.

Recently, several 3D cell shape representations have shown promising results. One approach uses spherical harmonics, which are a set of periodic functions defined on the surface of a sphere that form an orthonormal basis, a mathematical property that allows coordinates to be efficiently described by independent vectors. In this approach, the surface of each cell is mapped to a sphere, and the spherical harmonics transform is used to generate a list of coefficients that describe the cell’s shape.^9^ Principal components analysis (PCA) can then be applied to reduce dimensionality and facilitate interpretation of cell shape variation within and between cell populations. This method was recently used to understand variation in cell and nuclear shape of induced pluripotent stem cells in culture.^10^ Spherical harmonics have also been used to describe organ shape trajectories in murine limb and heart development,^11^ and have been used to describe 3D shapes in medical imaging.^12^ A strength of this method is that it maintains enough information to accurately reconstruct 3D cell shapes, enabling sophisticated, data-driven analysis of cell shape. Such a method, which eschews the limitations of choosing descriptive features or relying on 2D projections, could provide deeper insights into the relationships between cell shape, fate specification, and developmental trajectories. Here we test the idea that we can use these coefficients to cluster and classify cell types.

The lateral line system of zebrafish is an ideal model for studying cell shape in a living, developing animal. The lateral line is a sensory system that detects changes in water flow. Organs of the lateral line, known as neuromasts, are small, tightly packed epithelial rosettes with apical-basal polarity, radially organized cell types, and mirror symmetry.^13,14^ Mechanosensory hair cells are located in the center of the neuromast and surrounded by nonsensory, glia-like support cells. Support cells can be further divided into subpopulations based on their locations, gene expression patterns, and propensity to act as progenitors during regeneration.^15,16^ Known support cell subpopulations include peripheral cells (also known as mantle cells), dorsoventral cells, and anteroposterior cells, which can be identified in live animals using established knock-in lines.^16^ In addition, the superficial location and optical transparency of zebrafish larvae facilitate high resolution live imaging of neuromasts. Together with the aforementioned knock-in lines, this model system enables interrogation of the relationship between cell shape, location, and fate in neuromasts.

In this study, we developed a workflow to segment neuromast cells and nuclei and analyze their 3D shapes using spherical harmonics and PCA. We were able to model cell and nucleus shape using just 8 cell shape and 4 nucleus shape principal components (PCs). We found that hair cells were clearly distinct from support cells in our shape space, while support cells appeared to vary continuously in shape. Unsupervised clustering in cell shape PC space revealed that markers for support cell subpopulations were enriched within certain clusters, suggesting these cell types have distinct cell shape features. Analysis of mutants in *atoh1a*, encoding a conserved transcription factor necessary for hair cell differentiation,^17,18^ revealed the loss of the hair cell shape phenotype and expansion of other cell shape phenotypes. We also successfully built classifiers to predict hair cell identity from cell and nucleus shape features. Our work demonstrates that cell shape parameters can be used to characterize and classify cell types in neuromasts, laying the foundation for future studies of form, fate, and function in sensory epithelia.

## Results

### A Semi-automated Workflow Accurately Segments Zebrafish Lateral Line Cells and Nuclei

Neuromasts are small, epithelial organs with tightly packed, apically constricted cells; this structure presents challenges for imaging and segmentation (Figure 1A). For all experiments, we used the zebrafish *Tg(cldnb:LY-EGFP)* transgenic line that labels neuromast cell membranes.^19^ In addition, some zebrafish contained a knock-in allele that labeled a cell population of interest. We used fish heterozygous for *sost^w^*^215*Tg*^ or *sfrp1a^w^*^217*Tg*^ (henceforth referred to as *sost:NLS-Eos* and *sfrp1a:NLS-Eos*), which express nuclear-localized photoconvertible fluorescent protein Eos in dorsoventral support cells and peripheral support cells, respectively, and have no effect on neuromast development.^16^ Some zebrafish also contained one or two copies of *atoh1a^w^*^271*Tg*^, a targeted insertion of mRuby into the *atoh1a* locus that when homozygous results in loss of *atoh1a* function and loss of hair cells. For the rest of this paper, we refer to this knock-in line as *atoh1a:mRuby* and indicate whether heterozygous or mutant animals are being described. Additional analyses of these lines are described in later sections. For numbers of neuromasts and cells from each line, refer to Supplemental Table 1. All fish were stained with the far-red nuclear dye DRAQ5.^20^ To achieve sufficient spatial resolution, we used a confocal microscope equipped with AiryScan.^21^ Fish were imaged live at 5 days post fertilization (dpf), at which point neuromasts are functionally mature. Minimal cell death was observed during imaging; any neuromasts exhibiting significant amounts of cell death were excluded from further analysis.

**Figure 1:**
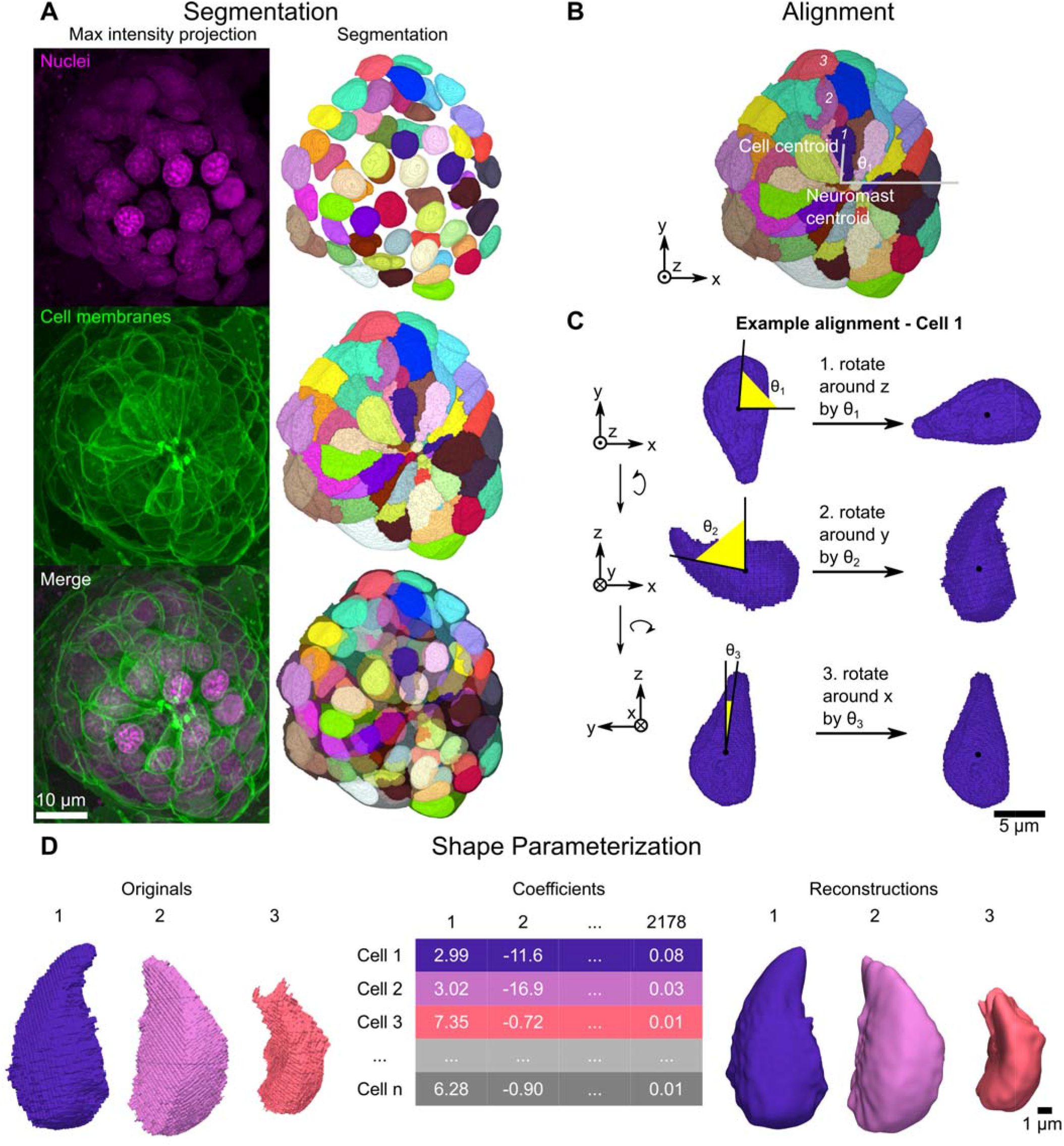
Overview of segmentation and shape parameterization. (A) Representative maximum intensity projections (left column) and corresponding 3D projections of segmentations (right column) for nuclei labeled by DRAQ5 dye (top row), cells labeled by *Tg(-8.0cldnb:LY-EGFP)* (middle row), and both channels together (bottom row). (B) Image depicting how the first rotation angle is calculated. Example given for the maximum intensity projection segmentation channel of the neuromast shown in (A). Lines extending from the neuromast centroid to the cell centroid and depicting the x-axis originating from the neuromast centroid are overlaid on the image. Cells 1, 2, and 3 used in (C) and (D) are labeled with the corresponding numbers. (C) Example of the alignment procedure as applied to cell 1 from (B). To correct for radial symmetry, cells are first rotated around the z axis. Cells were then rotated around the y axis to correct for apicobasal tilt, and then around the x axis to correct for off-parallel tilt during imaging. (D) Overview of shape parameterization using spherical harmonics, as applied to cells 1, 2, and 3 from (B). Spherical harmonics expansion (SHE) is applied to 3D surface meshes of the original, segmented cell surfaces (left) to generate the cell-by-SHE coefficients table (middle) which can then be used to generate 3D reconstructions (right) of each cell.

Due to the tight packing of neuromast cells and variations in GFP signal intensity, classical thresholding-based segmentation approaches did not yield accurate results for cells or nuclei in our images. To improve the accuracy of segmentation, we trained machine learning models to detect cell boundaries from the membrane GFP signal and nuclei masks from the DRAQ5 signal using the Allen Cell and Structure Segmenter.^22^ We then used the model results as the foundation for watershed-based instance segmentation. In essence, the watershed transform treats the image as a topographical map and identifies “catchment basins” as separate objects.^23^ We first applied a distance transform watershed algorithm to segment every nucleus. Labeled nuclei were then used as seeds for marker-based watershed of membrane boundary predictions. The accuracy of segmentation was further improved through proofreading and manual correction. This semi-automated workflow produced good quality segmentations for most cells (Figure 1A). After segmentation, each field of view was manually inspected, and some cells were excluded for quality control (e.g. due to having poor signal or extending beyond the edge of the field of view). 164 cells were manually excluded in this way. In total, 3274 cells from 49 different neuromasts across 6 different imaging sessions were segmented.

### Spherical Harmonics Accurately Represent Most Neuromast Cell Shapes

To quantitatively compare cell shapes, a list of numerical shape features must be extracted from each segmented cell in our dataset. In other words, the shape of each cell must be encoded as a vector. To approach this problem in an unbiased, data-driven fashion and maximize the amount of 3D shape information extracted from each cell, we used a method based on spherical harmonics expansion (SHE).^9,10^ In this method, the shape of the cell is mapped to the surface of a sphere, and the spherical harmonics transform is applied. The result is a vector of coefficients that describe the shape of the cell (organized into a cell-by-SHE-coefficients table), on which the inverse transform can be applied to create a reconstruction of the cell (Figure 1D).

One challenging aspect of this approach is that it is sensitive to the cell orientation – i.e., two cells with essentially the same shape that are rotated relative to each other will have different values of SHE coefficients. If differences in shape (rather than rotation) are of interest, then the cells must first be aligned to minimize rotational variation. We identified three sources of rotational variation in our dataset: (1) the radial symmetry of neuromast cells around the apical-basal axis of the rosette, (2) differential tilt of the cell along the radial axis as cells come together at the apical end of the rosette, and (3) the fact that some samples were slightly tilted during image acquisition such that the imaging z-axis was slightly off-parallel to the neuromast apical-basal axis. To correct for (1), cells were rotated around the imaging z-axis such that vectors pointing from the cell centroids to the neuromast centroid were aligned to the x-axis (and thus to each other). To correct for (2) and (3), two additional rotations were applied. The first aligned the long axis of the cell in the xz-plane to the z-axis. The second aligned the long axis of the cell in the yz-plane to the z-axis, with one caveat: if the rotation required would be more than 45 degrees, the long axis in the yz-plane was aligned to the y-axis instead. This adjustment in rotation angle accounts for the fact that some cells in the dataset were longer along the y-axis than the z-axis, and we wished to preserve their apical-basal orientation. An example of a cell aligned by this procedure is shown in Figure 1B,C.

Following alignment, we applied SHE to our dataset, generating 2178 spherical harmonics coefficients for each cell (Figure 1D). As a measure of reconstruction fidelity, we used the directed Hausdorff distance, a metric often used in image processing that is reflective of the largest error between the original and its reconstruction.^24^ We found a bimodal distribution of error, with 84% of cells accurately reconstructed (Supplemental Figure 1A). Cells that were not accurately reconstructed tended to be located midway between the center and periphery of the neuromast (Supplemental Figure 1B) and had shapes that rendered them unsuitable for spherical harmonics expansion, such as high concavity (Supplemental Figure 1C). Cells with high reconstruction error (421 cells, representing 16% of the original dataset) were excluded from further cell shape analysis. To further understand the types of shapes that were excluded, we measured additional properties of excluded cells. Compared to other cells, excluded cells tended to have greater cell surface area and height (Supplemental Figure 1D, top). Nuclei of excluded cells had similar dimensions to those of retained cells (Supplemental Figure 1D, bottom). In sum, some tall, curved cells located in a ring around the middle of the neuromast were not reconstructed well by spherical harmonics, and the shapes of these cells were not used in our subsequent analysis.

### Principal Components Analysis Reveals Variations in Cell Shape Between Neuromast Cell Types

To identify, visualize, and interpret the main axes of shape variation in the dataset, we applied principal components analysis (PCA) to the cell-by-SHE-coefficients matrix. The 2178 SHE coefficients were reduced to 8 principal components (PCs) representing approximately 70% of the total variance in cell shape (Supplemental Figure 2). A key advantage of this approach is that from any point in this 8-dimensional shape space, it is possible to calculate the corresponding SHE coefficients and create 3D representations of real or theoretical cells. Reconstructing cells in this way would not be possible if using a list of geometric features to describe cell shape.

Neuromasts contain several cell types with qualitatively different shapes. For example, lateral line hair cells have a round, flask-like morphology, while many support cells are concave in shape or have thin apical projections. We wondered whether these different morphologies would be reflected in our shape space, and if so, whether variation in shape between cell types would be continuous (indicating smooth transitions between different shapes) or discrete (suggesting some populations have features that set them more clearly apart from other cells). To visualize shape variation, each PC was first individually z-scored (i.e. the values of each PC were divided by that PC’s standard deviation) to generate “shape modes.” We used the point corresponding to the origin in shape space (where the value of all 8 cell shape modes is 0) to create a visual representation of the mean cell shape in the dataset. We then varied the value of one shape mode while keeping the values of the other shape modes constant to generate visual representations of each shape mode (Figure 2A). To compare the relationship between cell shape and position across neuromasts, we calculated the relative location of each cell using its radial distance and angular position (Figure 2B). A small number of interpretable parameters therefore captured the majority of cell shape variation, enabling quantitative comparisons of cell shapes within the dataset.

**Figure 2:**
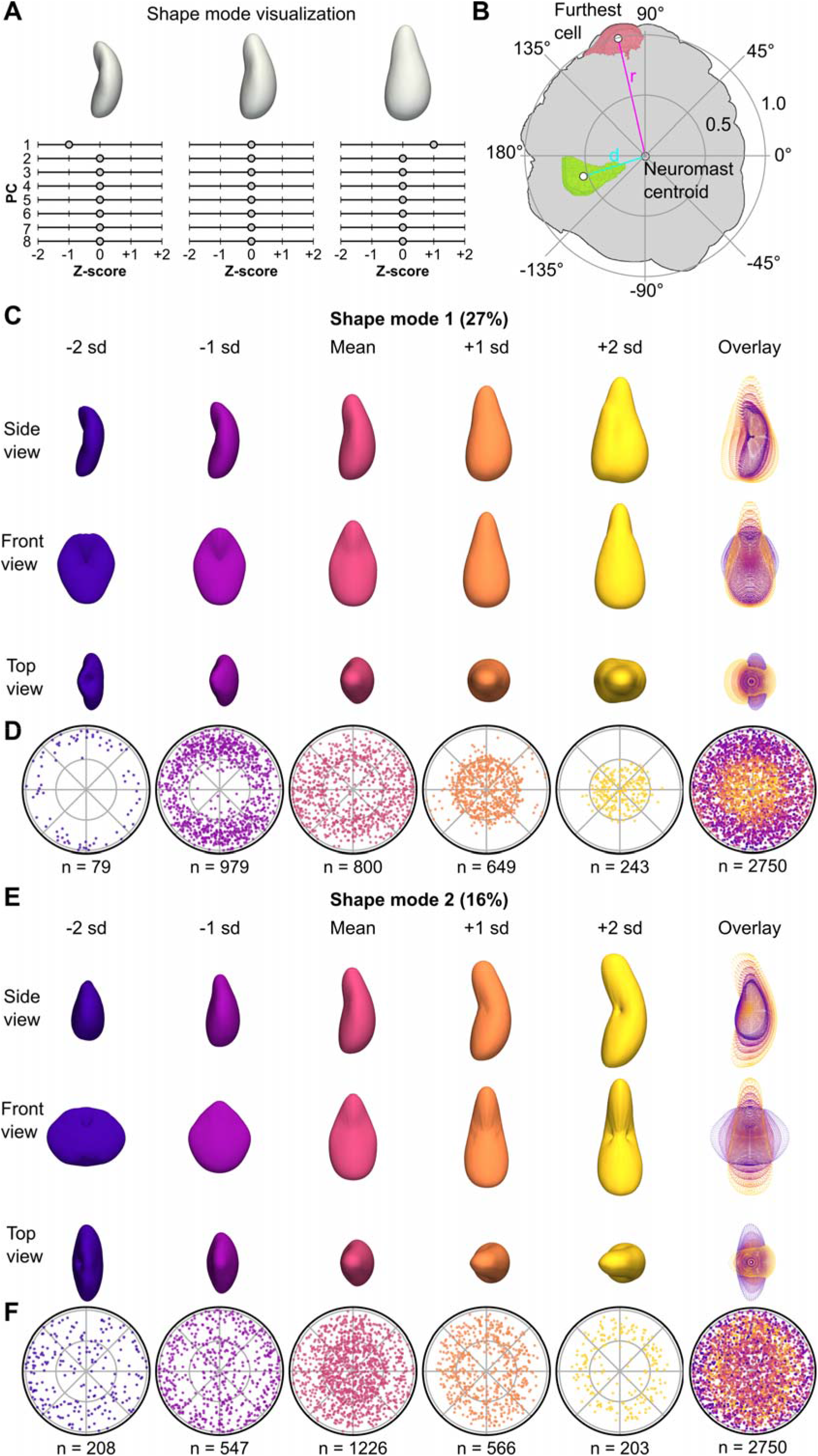
Overview of segmentation and shape parameterization. (A) Schematic depicting how shape variation was visualized. PCs were first z-scored to generate shape modes. Visual representations of each shape mode were generated by finding the mean cell (a hypothetical cell with a value of 0 for all 8 shape modes) and varying one shape mode (here, shape mode 1) by up to 2 standard deviations (sd) in increments of 1 sd while holding other shape modes at the mean (0). (B) Schematic depicting how relative cell locations within each neuromast were calculated. First, for each neuromast, the distance between the neuromast centroid and the furthest cell (r) in the XY plane was calculated. The distances between the neuromast centroid and cell centroids were divided by (r) to yield the normalized distance from the neuromast center (d) for each cell. The angular location of each cell was determined in reference to the positive x-axis centered on the neuromast centroid. Neuromast cell locations were thus projected onto a unit circle with radius 1, allowing relative cell locations from multiple neuromasts to be plotted together. (C) Visualization of the first mode of cell shape variation (Cell shape mode 1, CSM1), generated as described in (A). 3D surface renderings are shown from 3 different views, as well as an overlay of point representations for each view (rightmost column). (D) Polar plots of relative cell locations (generated as shown in B) binned by CSM1 scores. Bins were centered on the corresponding z-score and included cells with values within 0.5 (e.g., the second plot from the right corresponded to cells with CSM1 scores between -1.5 sd and -0.5 sd). For the -2 sd and 2 sd plots, the upper bounds were not limited to within 0.5 (e.g. the first plot includes all cells with CSM1 values < -1.5 sd). An overlay of the first 5 plots is shown in the rightmost column. (E) Visualization of the second mode of cell shape variation (Cell shape mode 2, CSM2), generated as described in (A). 3D surface renderings are shown from 3 different views, as well as an overlay of point representations for each view (rightmost column). (F) Polar plots of relative cell locations (generated as shown in B) binned by CSM2 scores. An overlay of the first 5 plots is shown in the rightmost column.

We hypothesized that hair cells, which can be identified by eye based on shape, would be quantitatively distinct from other cells using our shape analysis method. To test this hypothesis, we manually annotated hair cells in our dataset using the presence of apical stereocilia, which are visible in the *Tg(cldnb:LY-EGFP)* transgenic line. We found that cell shape mode 1 (CSM1, representing 27% of total variance) appeared to represent “hair cell-ness.” Positive CSM1 scores corresponded with greater cell volume and width (Supplemental Figure 3B), as well as a more convex, flask-like shape (Figure 2C). Cells with positive CSM1 scores also tended to be closer to the neuromast center (Figure 2D, Supplemental Figure 3D), consistent with the characteristic location of hair cells. This result suggests that much of cell shape variation in neuromasts is due to differences between hair cells and support cells.

In contrast to CSM1, cell shape mode 2 (CSM2, representing 16% of total variance) did not distinguish between hair cells and support cells; rather it appeared to distinguish among support cell types. Cells with increasing CSM2 scores were progressively taller and less deep (Figure 2E, Supplemental Figure 3B). CSM2 was also negatively associated with being located toward the neuromast periphery, particularly for support cells (Figure 2F, Supplemental Figure 3D). Hair cells tended to have average CSM2 scores, clustered around zero, and showed a narrower distribution than support cells (Supplemental Figure 3C). Taken together, the relationships between cell type and cell position likely explain the “bullseye” pattern observed when CSM2 scores are plotted by relative cell location: support cells have higher CSM2 scores as they become more central, while hair cells (which are also centrally located) have CSM2 scores close to zero (Figure 2F).These observations suggest that CSM2 represents shape variation between support cells in a way that is distinct from the variation between support cells and hair cells.

To facilitate interpretation of shape modes and further query the relationships between shape modes and cell type, we compared shape modes to geometric features and location parameters (Supplemental Figure 3). We correlated CSM1 and CSM2 to cell width, depth, height, volume, surface area and cell distance from the neuromast center (Supplemental Figure 3B,D). We found a positive correlation between CSM1 and cell volume and cell width, and a negative correlation with distance from the center of the neuromast. For CSM2, there was a positive correlation with cell height and a negative correlation with cell depth and distance from center. Hair cells consistently segregated with CSM1 but not CSM2 (Supplemental Figure 3C,D). Beyond CSM1 and CSM2, other shape modes appeared to represent more subtle ways in which neuromast cell shape varied (Supplemental Figure 2).

### Nuclear Shape Variation Is Captured by SHE and PCA

We also performed SHE and PCA on nucleus shape. Nuclei were aligned to account for position within the neuromast using a series of rotations similar to (but independent of) that used to align cells (see Methods for details). Because nuclei had a much lower reconstruction error compared to cells (Supplemental Figure 1A), we elected not to exclude any nuclei from analysis and maintained the original dataset of 3274 cells to see whether additional insights about these cells could be gained from nuclear shape analysis. Nuclear SHE coefficients were reduced to 4 principal components (PCs) representing approximately 76% of the total variance in nuclear shape (Supplemental Figure 4). As for cell shape analysis, nuclear shape PCs were z-scored to describe them as nuclear shape modes.

Nucleus shape mode 1 (NSM1, representing 40% of total variance, Figure 3A) and nucleus shape mode 2 (NSM2, representing 18% of total variance, Figure 3C) captured distinct aspects of variation in nuclear shape. Increasing NSM1 scores were associated with a shift from short, wide shapes to a tall, crescent-like morphology (Figure 3A). Meanwhile, NSM2 appeared to describe a progression from tall and cylindrical to flattened, rectangular shapes (Figure 3C). Nucleus shape modes 3 and 4 are summarized in Supplemental Figure 4. Comparing to other nuclear shape parameters (Supplemental Figure 5), we found NSM1 was negatively correlated with nucleus width and positively correlated with nucleus height (Supplemental Figure 5B,D). NSM2 was positively correlated with nucleus depth and negatively correlated with nucleus volume (Supplemental Figure 5B,D).

**Figure 3:**
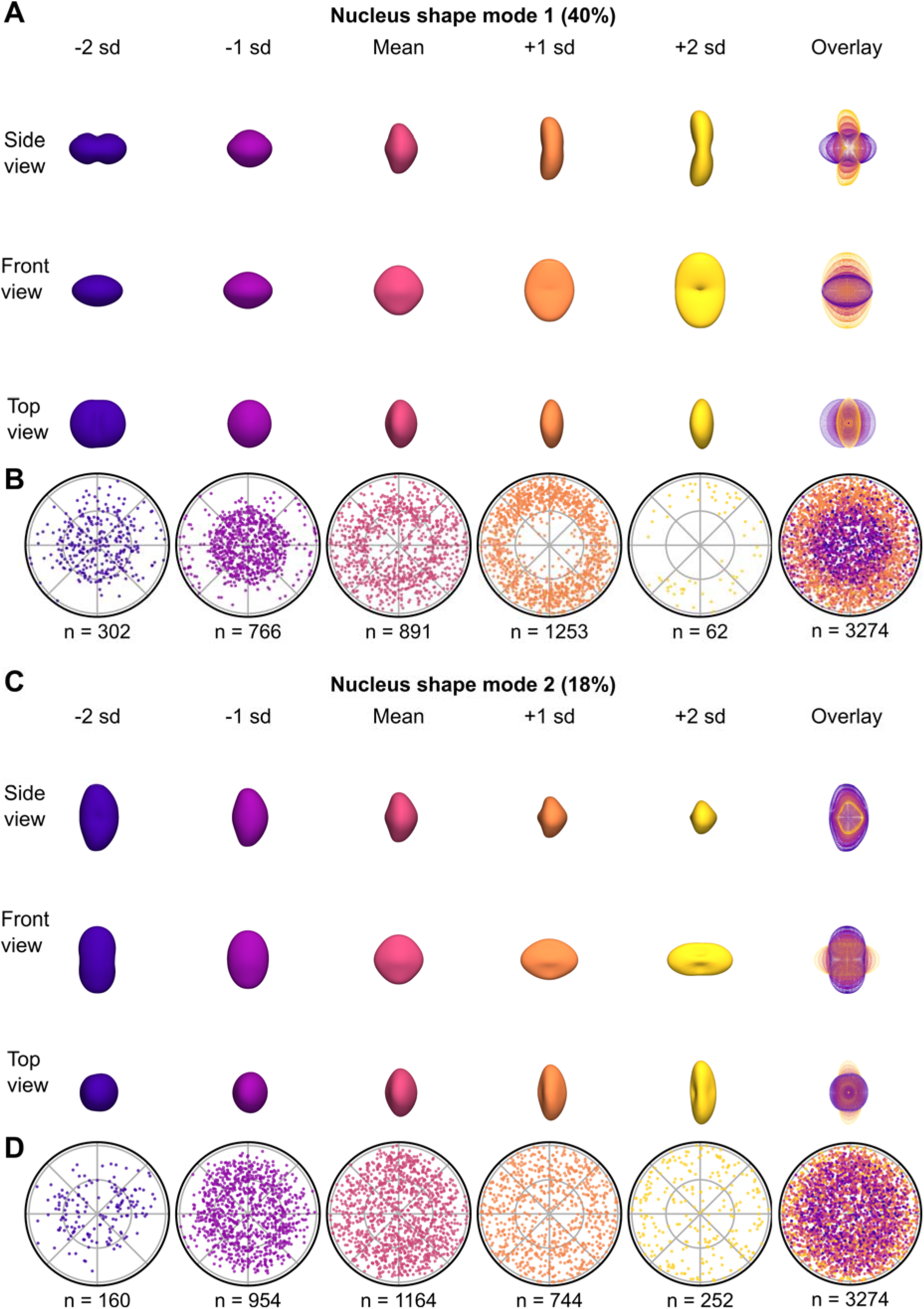
Overview of segmentation and shape parameterization. (A) Visualization of the first mode of nucleus shape variation (Nucleus shape mode 1, NSM1), generated as described in Figure 2A. 3D surface renderings are shown from 3 different views, as well as an overlay of point representations for each view (rightmost column). (B) Polar plots of relative cell locations (generated as shown in Figure 2B) binned by NSM1 scores. Bins were centered on the corresponding z-score and included cells with values within 0.5 (e.g., the second plot from the right corresponded to cells with NSM1 scores between -1.5 sd and -0.5 sd). For the -2 sd and 2 sd plots, the upper bounds were not limited to within 0.5 (e.g. the first plot includes all cells with NSM1 values < -1.5 sd). An overlay of the first 5 plots is shown in the rightmost column. (C) Visualization of the second mode of nucleus shape variation (Nucleus shape mode 2, NSM2), generated as described in Figure 2A. 3D surface renderings are shown from 3 different views, as well as an overlay of point representations for each view (rightmost column). (D) Polar plots of relative cell locations (generated as shown in Figure 2B) binned by NSM2 scores. An overlay of the first 5 plots is shown in the rightmost column.

We next asked to what degree nuclear shape modes distinguished among hair cells and support cells. We found that NSM1 had a bimodal distribution (Supplemental Figure 5C) and was correlated with distance from the neuromast center (Figure 3B, Supplemental Figure 5D). Like CSM1, NSM1 also appeared to distinguish between hair cells and support cells, but to a lesser degree since some support cells also had high NSM1 scores (Supplemental Figure 5C). NSM2 also showed a slightly skewed distribution and a correlation with distance to the neuromast center (Figure 3D, Supplemental Figure 5D), with some distinction between hair cells and support cells. Nuclear shape modes therefore also captured variation between hair cells and support cells, although this information was split across multiple shape modes.

### Unsupervised Clustering of Neuromast Cells in Shape Space Identifies Groups of Cells With Similar Shapes and Locations

We next wanted to determine whether shape features could be used to assign neuromast cells to biologically meaningful categories beyond hair cells and support cells. To do so, we performed unsupervised clustering on the first 8 cell shape PCs using the Phenograph package with the Leiden community detection algorithm.^25,26^ Five clusters were identified and represented in two dimensions by a uniform manifold approximation and projection (UMAP, Figure 4A)^27^ where each cell is a point colored by cluster label. The fraction of cells within each cluster was proportional across all individual neuromasts (Supplemental Figure 7D) suggesting that clustering is influenced by shared shape parameters and not due to differences between neuromasts. To quantify the relationships between clusters, we performed partition-based graph abstraction (PAGA).^28^ Relative strength of association is represented by line thickness (Figure 4B). PAGA analysis revealed relatively greater similarities between clusters 0/2, 1/2/3, and 2/4.

**Figure 4:**
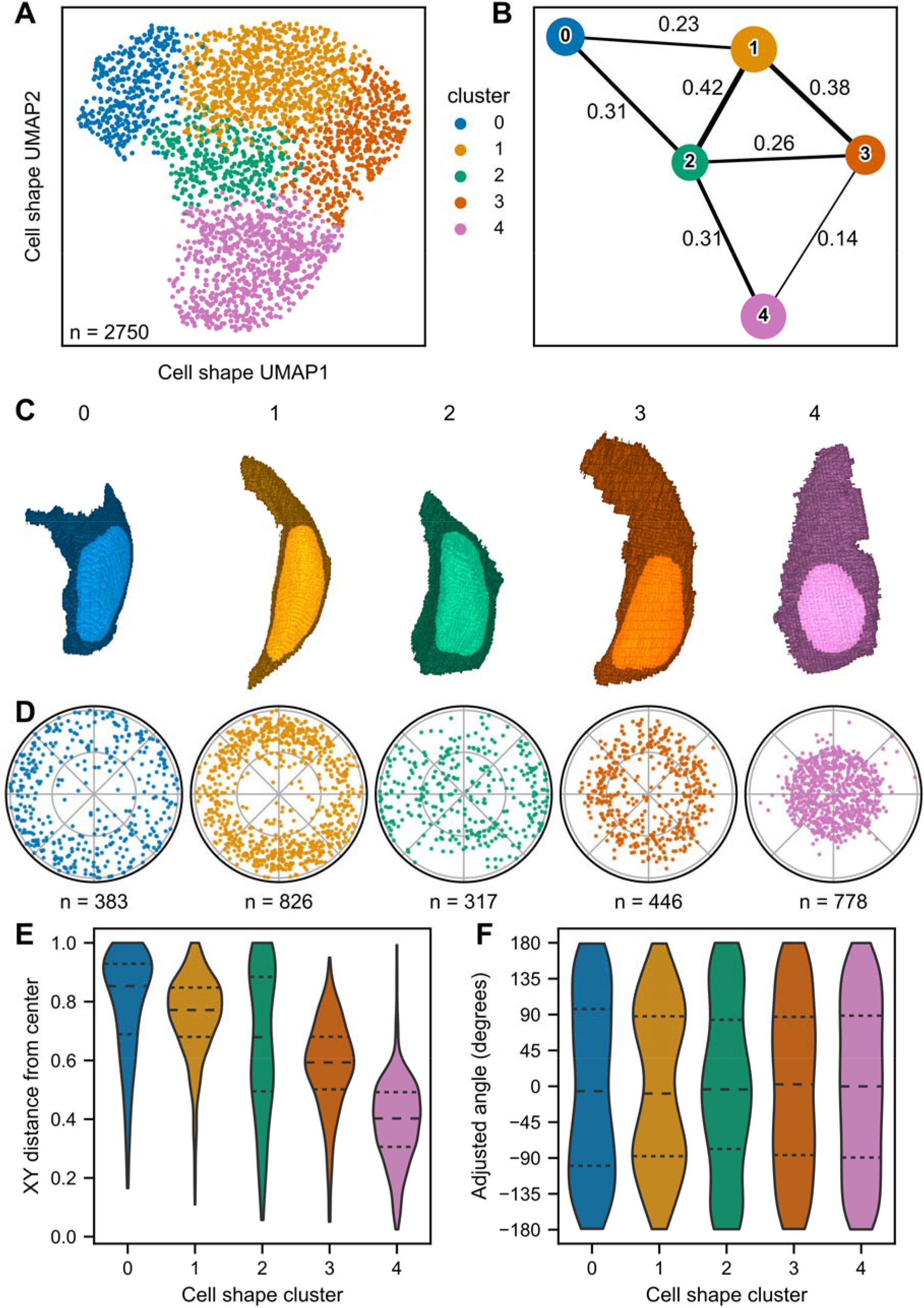
Overview of segmentation and shape parameterization. (A) UMAP of neuromast cells, color coded by cell shape cluster. Colors and associated cluster numbers are used in (B), (C), (D), (E), and (F). (B) PAGA plot of cell shape clusters. Each node represents a cluster. Thickness of the line drawn between nodes indicates estimated connectivity between clusters (the value of which is shown adjacent to the corresponding line). Edges with connectivity below 0.1 are not depicted. (C) 3D projections of representative cells for each cluster, defined as the cell closest to the cluster centroid in eight-dimensional PC space. The original segmentation masks (not SHE reconstructions) are depicted. (D) Polar plots of relative neuromast locations of cells within each cluster. (E) Distributions of the cell distance from the neuromast center for each cluster. Dashed lines indicate quartiles. Distances were calculated as the distance between the xy coordinates of cell centroids and the xy coordinates of the corresponding neuromast centroid. Distances were normalized to that of the cell with the greatest distance from the center (see Figure 2B). (F) Distributions of neuromast cell angles for each cluster. Dashed lines indicate quartiles. Angles were calculated with reference to the positive x-axis extending from the neuromast centroid. Cluster 1 shows a bias towards the dorsoventral poles (90, -90), while cluster 2 shows a bias to the anterior-posterior compartments (0, 180, -180).

To visualize the characteristic shapes of cells within each cluster and facilitate comparisons between them, we identified the cell closest to the centroid of each cluster in eight-dimensional PC space as a “representative cell” (Figure 4C). Cells in clusters 0 tended to be shorter with greater depth, while cells in cluster 3 tended to be tall and wide. Cells in clusters 1 and 2 appeared to have shapes intermediate between clusters 0 and 3. Meanwhile, cells in cluster 4 tended to have flask shapes reminiscent of hair cells; quantifying manually annotated hair cells in each cluster confirmed that most hair cells were assigned to cluster 4 (Supplemental Figure 7E) and cluster 4 was primarily composed of hair cells (Supplemental Figure 7F).

Cells from a given cluster tended to be found in similar locations across neuromasts in the dataset. In particular, cells in specific clusters were found at distinct radial distances from the neuromast center (Figure 4D,E), and we used this order to assign cluster number labels. However, we also found that some clusters were distributed in distinct quadrants of the neuromast (Figure 4D,F). This difference in distribution was particularly apparent for cluster 1, where cells were more concentrated within the dorsoventral quadrants of the neuromast (Figure 4F, represented by adjusted angle 90 and -90), and cluster 2, where cells were more often found within the anteroposterior quadrants (Figure 4F, represented by adjusted angle 0). These results suggest that grouping neuromast cells purely using shape descriptors, without explicitly including information about cell type or identity, recapitulates known aspects of neuromast organization (e.g. concentric and radially symmetric cell populations).

We also performed unsupervised clustering on the first 4 nucleus shape PCs. Similar to cell shape clusters, we found distinct nucleus shape clusters (Figure 5A) whose relative relationships could be measured through PAGA analysis (Figure 5B). As for cells, we identified representative nuclei for each cluster (Figure 5C). Clusters 4 and 5 were primarily composed of hair cells, and cluster 1 was enriched for dorsoventral cells (Figure 5D). Nucleus shape clusters exhibited distinct spatial locations within the neuromast (Figure 5D,E,F). However, nucleus shape clusters did not appear to distinguish as well between support cell subpopulations compared to cell shape clusters. Since the nucleus shape dataset contains cells that were excluded from cell shape analysis, we looked at the distribution of excluded cells across nucleus shape clusters (as a way to understand whether a particular class of cells had been excluded). We found excluded cells in all 5 nucleus shape clusters, suggesting that excluded cells can have a range of nucleus shape phenotypes (Supplemental Figure 1E). Based on these results, we decided to perform the remaining analysis using only cell shape clusters.

**Figure 5:**
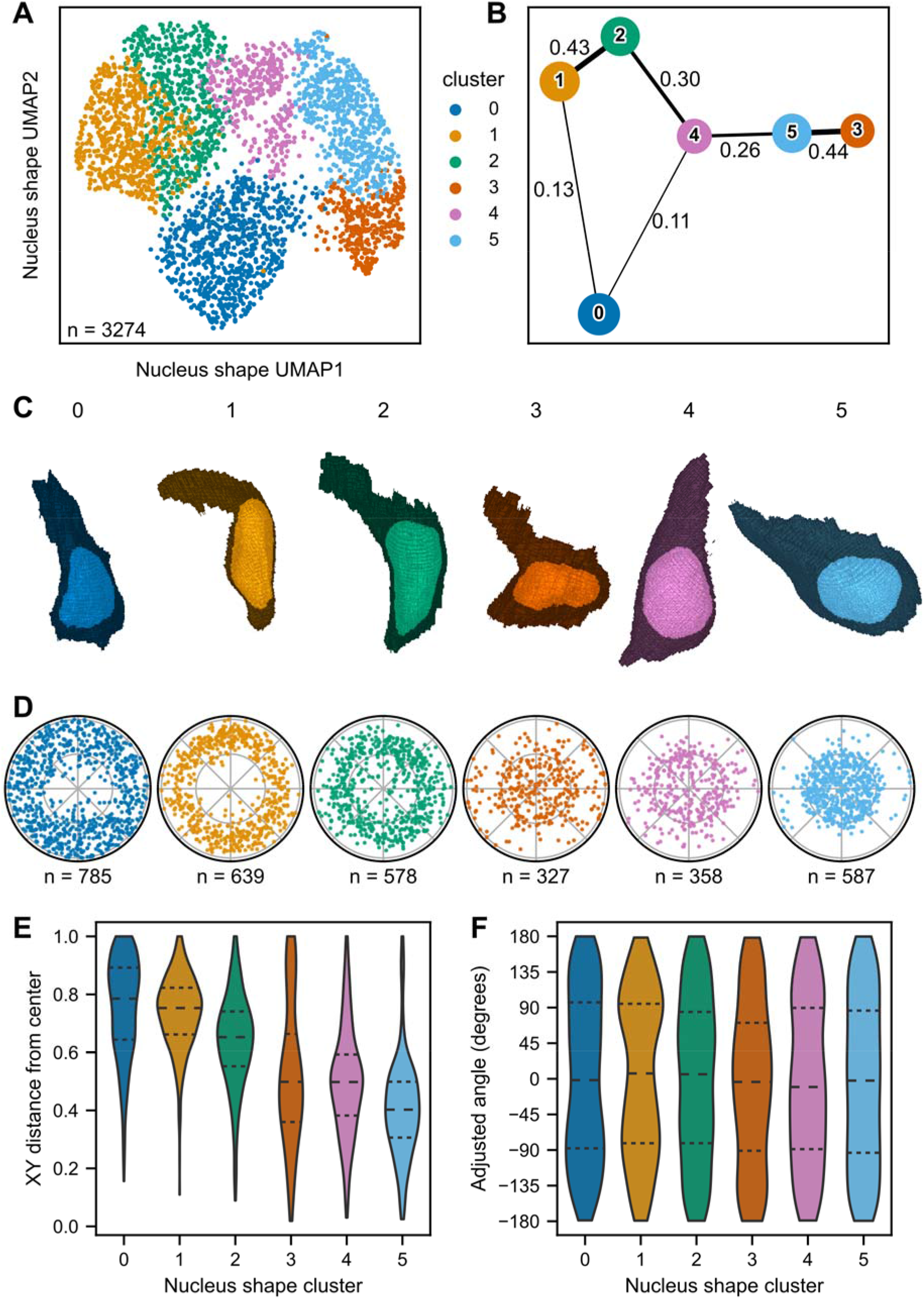
Overview of segmentation and shape parameterization. (A) UMAP of neuromast cells, color coded by nucleus shape cluster. Colors and associated cluster numbers are used in (B), (C), (D), (E), and (F). (B) PAGA plot of nucleus shape clusters. Each node represents a cluster. Thickness of the line drawn between nodes indicates estimated connectivity between clusters (the value of which is shown adjacent to the corresponding line). Edges with connectivity below 0.1 are not depicted. (C) 3D projections of representative cells for each nuclear cluster, defined as the cell closest to the cluster centroid in four-dimensional PC space. The original segmentation masks are depicted (not SHE reconstructions). (D) Polar plots of relative locations of cells within each cluster. (E) Distributions of the cell distance from the neuromast center for each cluster. Dashed lines indicate quartiles. Distances were calculated as the distance between the xy coordinates of cell centroids and the xy coordinates of the corresponding neuromast centroid. Distances were normalized to that of the cell with the greatest distance from the center. (F) Distributions of neuromast cell angles for each cluster. Dashed lines indicate quartiles. Angles were calculated with reference to the positive x-axis extending from the neuromast centroid.

### Cell Fate Markers of Support Cell Subpopulations Are Associated With Distinct Cell Shape Phenotypes

Clustering purely based on shape descriptors yielded groups with distinct spatial patterns in neuromasts resembling those of previously known support cell types. For example, peripheral support cells (mantle cells) form a ring around the outside of the neuromast. Distinct support cell subpopulations also exist within the dorsoventral (DV) and anteroposterior (AP) compartments of the neuromast. These support cell subpopulations have characteristic transcriptional profiles and behaviors during regeneration:^15,16^ DV cells act as hair cell progenitors while peripheral cells can replenish DV cells depleted by hair cell differentiation. We hypothesized that if cell shape clusters correspond to cells with distinct identities, they would be enriched for markers of these support cell subpopulations.

To examine the relationship between support cell identity and cell shape, we examined neuromasts from knock-in lines that mark distinct support cell subpopulations. The numbers of neuromasts and cells analyzed for each line are summarized in Supplemental Table 1. The *sost:NLS-Eos* line marks cells in the DV compartment of the neuromast (Figure 6A-C). We measured the mean intensities of Eos fluorescence in each cell, normalizing to z-score for each individual neuromast. We then classified cells as Eos+ if they had fluorescence with z-score greater than 1 (Figure 6B, Supplemental Figure 6A). An example neuromast with labeled positive cells is shown in Figure 6B. As expected, *sost:NLS-Eos*+ cells were found in DV quadrants (Figure 6B,C). We next asked whether *sost:NLS-Eos*+ cells were differentially distributed across clusters (Figure 6E,F). Most *sost:NLS-Eos+* cells were found in cluster 1 (Figure 6E). 33% of cluster 1 cells were s*ost:NLS-Eos*+, the highest proportion of any cluster (Figure 6F). A small number of *sost:NLS-Eos+* cells were also found in clusters 2 and 3,making up 13% and 17% of clusters 2 and 3, respectively (Figure 6F). These results indicate that DV support cells, as measured by *sost:NLS-Eos* expression, occupy a subset of possible cell shape states within neuromast cells.

**Figure 6:**
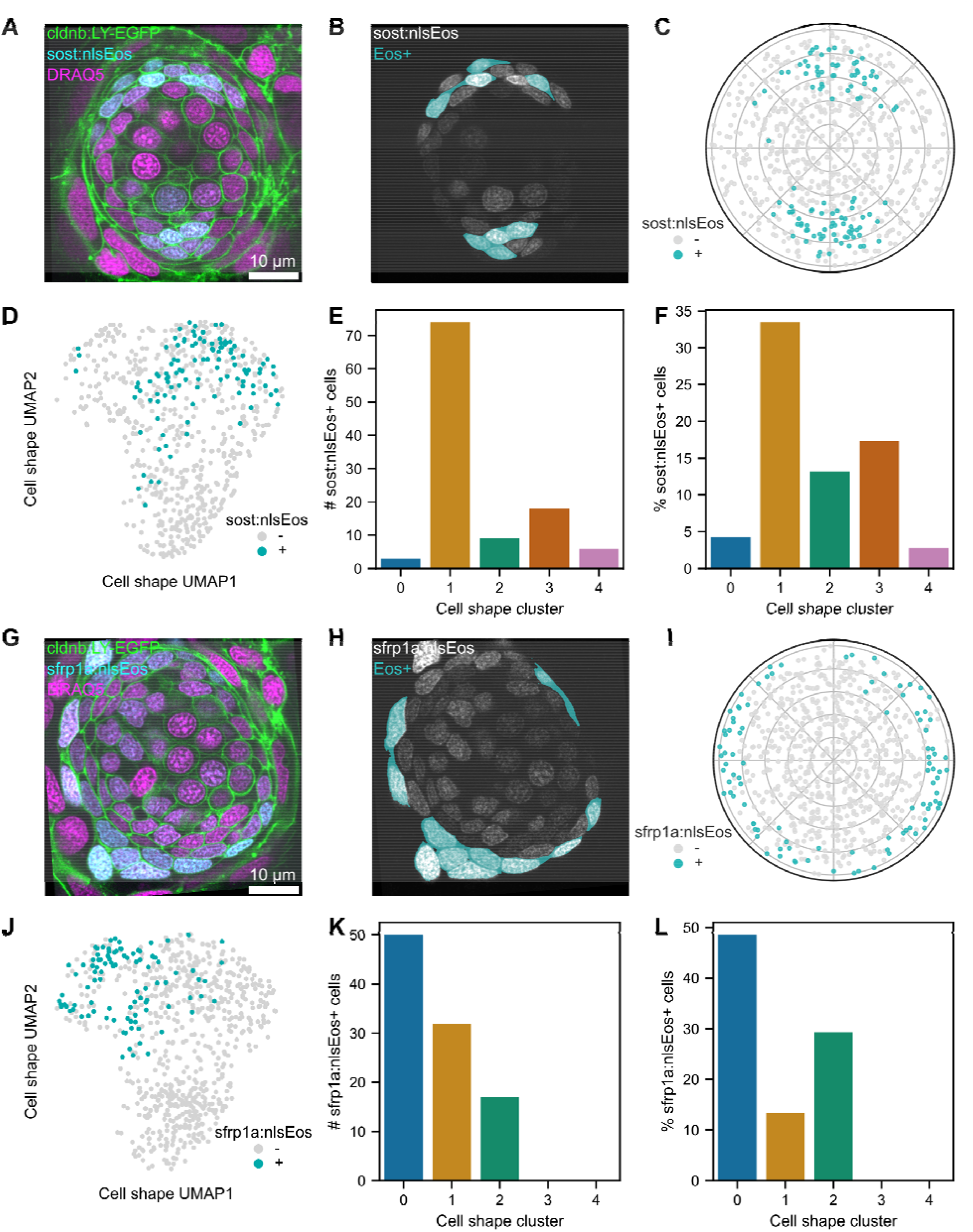
Location and shape characteristics of cells expressing *sost:NLS-Eos* and *sfrp1a:NLS-Eos*. (A) Single confocal slice of a neuromast from *sost:NLS-Eos; Tg(-8.0cldnb:LY-EGFP)* fish stained with DRAQ5. Image contrast was adjusted for visibility. (B) The same slice as in (A), with the *sost:NLS-Eos* channel (gray) overlaid with a mask indicating cells classified as Eos+ (dark cyan). (C) Polar plot showing relative cell locations of locations of Eos+ (dark cyan) and Eos-(gray) cells. (D) UMAP of Eos+ (dark cyan) and Eos-(gray) cells from *sost:NLS-Eos*. (E) *sost:NLS-Eos+* cells plotted as raw counts within each cell shape cluster. (F) *sost:NLS-Eos+* cells plotted as the percentage of each cell shape cluster. (G) Single confocal slice of a neuromast from *sfrp1a:NLS-Eos; Tg(-8.0cldnb:LY-EGFP)* fish stained with DRAQ5. Image contrast was adjusted for visibility. (H) The same slice as in (A), with the *sfrp1a:NLS-Eos* channel (gray) overlaid with a mask indicating cells classified as Eos+ (dark cyan). (I) Polar plot showing relative cell locations of locations of Eos+ (dark cyan) and Eos-(gray) cells. (J) UMAP of Eos+ (dark cyan) and Eos-(gray) cells from *sfrp1a:NLS-Eos*. (K) *sfrp1a:NLS-Eos+* cells plotted as raw counts within each cell shape cluster. (L) *sfrp1a:NLS-Eos+* cells plotted as the percentage of each cell shape cluster.

The *sfrp1a:NLS-Eos* line marks peripheral support cells (Figure 6G-I). We again measured the mean intensities of Eos fluorescence in each cell, normalizing to z-score, and classified cells as Eos+ with fluorescence with z-score greater than 1 (Figure 6H, Supplemental Figure 6B). Cells expressing higher levels of Eos were found in the neuromast periphery (Figure 6H,I). Most *sfrp1a:NLS-Eos+* cells were found within cluster 0, making up about half of the cells in this cluster (Figure 6K,L). Some *sfrp1a:NLS-Eos+* cells were also found in clusters 1 and 3, making up 13% and 29% of clusters 1 and 3, respectively. As with DV support cells, peripheral support cells tended to inhabit a smaller range of shapes within the neuromast cell shape space. Overall, there was no difference in the distribution of clusters among individual neuromasts across wildtype, *sost:NLS-Eos*, and *sfrp1a:NLS-Eos* lines (Supplemental Figure 6D). Taken together, these results demonstrate that cells with distinct identities, as indicated by *sost:NLS-Eos* and *sfrp1a:NLS-Eos* expression, can be recognized by distinct cell shape characteristics.

### *atoh1a* Mutants Exhibit Changes in Cell Shape Across Cell Populations

We next sought to test what happens to cell shape distributions when a single cell type is absent. The transcription factor *atoh1a* is required for hair cell fate specification in zebrafish lateral line neuromasts.^18,29^ We expected that the loss of the hair cell shape phenotype would be readily observable in *atoh1a* mutants. However, we wondered whether support cells would maintain the same shapes seen in wildtype fish or if their shapes would also be altered in *atoh1a* mutants. In the *atoh1a:mRuby* line, the mRuby fluorophore was inserted into the coding sequence of the *atoh1a* gene, disrupting its function but acting as a readout for *atoh1a* promoter activity. This knock-in line enables comparisons not only between wildtype, mutant, and heterozygous cell populations as a whole, but also between mRuby+ and mRuby-populations in each genotype.

In heterozygous fish, mRuby+ cells are localized to the center of the neuromast (Figure 7A; Supplemental Figure 6E). We measured the mean intensities of mRuby fluorescence in each cell, normalizing to z-score, and classified cells as mRuby+ if they had fluorescence with z-score greater than 1 (Figure 7B, Supplemental Figure 6C). Overall, the proportions of cells in each cluster were similar to those of wildtype fish (Supplemental Figure 6D,G). However, mRuby+ cells were differentially proportioned across clusters (Figure 7C,D). We found that mRuby+ cells were highly enriched in cluster 4 (Figure 7D). We confirmed that cluster 4 represented hair cells by examining the distribution of manually annotated hair cells in our dataset identified by the presence of apical stereocilia (Supplemental Figure 6H,I).

**Figure 7:**
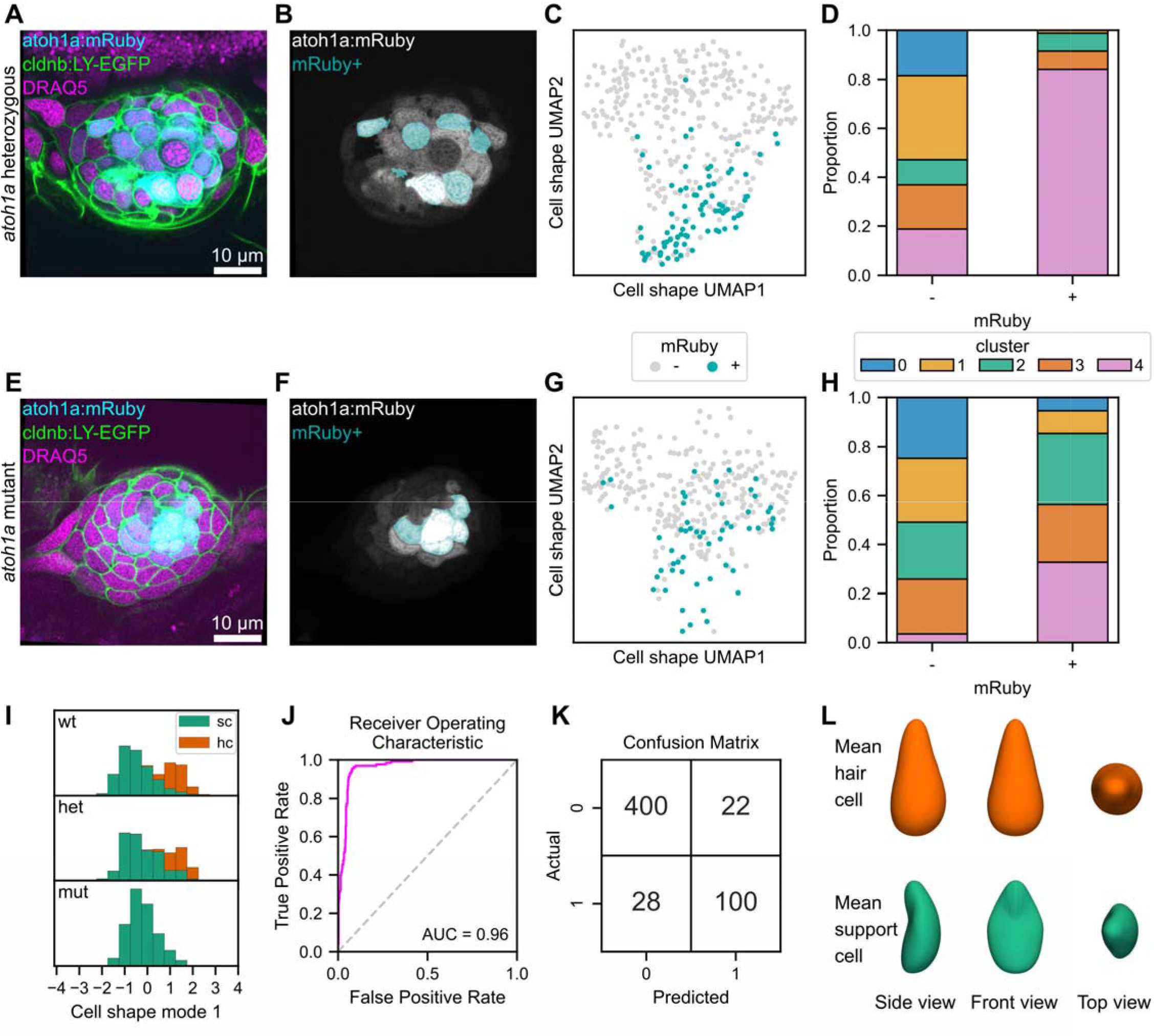
Shape characteristics of *atoh1a:mRuby* heterozygotes and mutants and logistic classifier for hair cells based on cell shape features. Analysis of cell shape distributions in *atoh1a:mRuby* heterozygotes (A-D) and mutants (E-G). (A, E) Single confocal slice of a neuromast from a *atoh1a:mRuby* heterozygote (A) and mutant (E) also expressing *Tg(-8.0cldnb:LY-EGFP)* and stained with DRAQ5. Image contrast was adjusted for visibility. (B, F) The same slices as in (A) and (E), with the *atoh1a:mRuby* channel (gray) overlaid with a mask indicating cells classified as mRuby+ (dark cyan) for a *atoh1a:mRuby* heterozygote (B) and mutant (F). (C, G) UMAP of mRuby+ (dark cyan) and mRuby-(gray) cells from *atoh1a:mRuby* heterozygotes (C) and mutants (F). (D, H) Proportions of mRuby-cells (left bar) and mRuby+ cells (right bar) from *atoh1a:mRuby* heterozygotes (D) and mutants (H) in each cell shape cluster. (I) Distributions of cell shape mode 1 scores, separated by genotype (top row: wildtype; middle row: *atoh1a* heterozygous; bottom row: *atoh1a* mutant). Hair cells (hc, green) and support cells (sc, orange) are color coded and displayed as stacked histograms. (J) Receiver Operating Characteristic (ROC) curve (fuchsia) for a logistic regression classifier trained to detect hair cells based on cell shape mode 1-8 scores. The Area Under the Curve (AUC) score is shown in the lower left of the plot (0.96). The silver dashed line on the diagonal represents the performance of a random classifier. (K) Confusion matrix for the cell shape-based hair cell classifier. 0 = support cells; 1 = hair cells. Top left: true negatives; top right: false positives; bottom left: false negatives; bottom right: true positives. (L) The idealized mean cell shape for hair cells (top, orange) and support cells (bottom, green) within the dataset.

In homozygous mutant fish, mRuby+ cells were still found in the center of the neuromast even though they did not differentiate into hair cells (Figure 7E,F; Supplemental Figure 6E). There was not a statistically significant difference in the proportion of mRuby+ cells between heterozygous and homozygous animals (Χ^2^ = 1.8, df = 1, p = 0.18; Supplemental Figure 6F). Like in heterozygotes, mRuby+ cells in homozygotes were differentially distributed across clusters (Figure 7G,H; Supplemental Figure 7D,G). Relative to wildtype and heterozygotes, mutant fish had far fewer cells in cluster 4 and an increased proportion in cluster 2 (Figure 7H; Supplemental Figure 7D,G). In heterozygotes, 84% of mRuby+ cells were found in cluster 4 (Figure 7D), but only 33% of mRuby+ cells were assigned to cluster 4 in mutants (Figure 7H). By contrast, the number of mRuby+ cells in clusters 2 and 3 were expanded in mutants. For mRuby-cells, cluster 2 was expanded in mutants relative to heterozygotes (Figure 7D,H). These results suggest that loss of *atoh1a* function is associated with a loss of the hair cell shape phenotype, an effect mainly seen in mRuby+ cells, and a concurrent shift in support cell shapes towards a greater number of cluster 2 phenotypes. The loss of hair cells did not result in the appearance of a new cell shape cluster but rather assignment of cells into existing shape clusters. These results demonstrate that cell shape analysis can be used to query changes in phenotype in contexts where development has been perturbed. Additionally, these results show that some aspects of cell shape are genetically determined in our system, rather than being solely due to other factors (such as position within the neuromast).

### Shape Modes Can Be Used to Classify Hair Cells

Our results thus far have suggested that hair cells have distinct cell shape phenotype compared to support cells. Previously, we observed that CSM1 scores appear to represent “hair cell-ness” (Figure 2C). A histogram of CSM1 scores exhibited a bimodal distribution with a positive and negative peak (Figure 2C), unlike the other shape modes that centered around zero (Supplemental Figure 2A). Comparing manually annotated hair cells to support cells, we found that hair cells tended to have positive CSM1 scores and comprised the majority of the positive peak in the shape mode histogram (Supplemental Figure 3C). To confirm this association, we examined the distribution of CSM1 scores of cells from *atoh1a* heterozygous and mutant fish. We found that the distribution of CSM1 scores in heterozygotes resembled that of wildtype fish, with hair cells associated with positive scores (Figure 7I, top and middle). By contrast, the distribution of CSM1 scores in homozygous mutants was no longer bimodal (Figure 7I, bottom) and was skewed negative and corresponded to the distribution of support cells in wildtype and heterozygous animals. CSM1 therefore captured the variation in shape between the two broadly defined neuromast cell populations (hair cells and support cells) and was predictably perturbed in mutants lacking hair cells.

The distinction between hair cells and support cells in CSM1 scores suggested that cell shape modes may contain sufficient information to classify cells as belonging to either of these cell types. We trained a logistic regression classifier to predict hair cell identity given the first eight cell shape mode scores (CSM1-8). The Receiver Operating Characteristic (ROC) curve indicated robust classifier performance with Area Under the Curve (AUC) of 0.96 (Figure 7J). Despite the imbalance of classes in the dataset (77% support cells and 23% hair cells), the classifier correctly labeled most (78%, 100/128) of the hair cells in the test set, while few support cells (5%, 22/422) were incorrectly labeled as hair cells (Figure 7K). We also asked if hair cells could be classified using nucleus shape features. We trained a logistic regression classifier using the first four nucleus shape modes (NSM1-4). Our classifier based on nucleus shape features performed similarly well to one using cell shape features (AUC score = .0.96; data not shown). Therefore, nuclear shape features can also be used to classify hair cells from support cells.

To visualize the shape characteristics associated with hair cell identity, we created a 3D reconstruction of the idealized mean hair cell and support cell in the dataset by calculating the mean CSM1-8 scores for each cell type and deriving the corresponding SHE coefficients (Figure 7L). As expected, the “average” hair cell has a flask-like morphology, while the “average” support cell has a concave shape. These results demonstrate the utility of cell shape features both for describing and classifying cell types.

## Discussion

During development and regeneration, cells transition through states as they grow, divide, and differentiate. Measuring the transcription of genes has proven to be a powerful method for understanding and classifying cell types. Using single cell RNA-sequencing, thousands of transcripts can be sequenced for each cell, generating high-dimensional data for cell type annotation and classification. However, cells can change in ways that are not necessarily reflected in RNA transcript levels.^30,31^ Cell shape analysis offers another avenue to quantitatively characterize and classify cells. Methods to encode cell shape in an unbiased, high dimensional representation have been employed with great success to cells in culture.^7,8,10^ However, to our knowledge, most studies of cell shape in living, developing animals have relied on 2D projections of cells or representative 3D geometric features, which may not capture many aspects of cell shape in these tissues. In this study, we use spherical harmonics and PCA to encode cell shape information in a more complete, unbiased fashion, allowing for data-driven analysis of the relationships between cell shape and fate.

A primary goal of our study was to understand the relationship between cell shape and established indicators of cell identity in neuromasts. If cell shape and markers for neuromast cell fate were tightly correlated, then it might be possible to classify cell types using shape features. Our results suggest that this relationship was true of hair cells, since we were able to build a logistic classifier for hair cells using our shape modes. In addition, when we performed unsupervised clustering of cells in shape space, there was a high degree of overlap between hair cell identity and one of the clusters (hair cells comprised 79% of cluster 4). Few hair cells were found in the other four clusters. In sum, using 3D cell shape analysis, we quantitatively captured differences between two broad classes of cells in our dataset. We believe a similar method could be used to build classifiers for cell types in other developmental contexts, potentially allowing for cell fate to be predicted from image-based data without expression of cell fate markers.

The association between cell shape and support cell subtypes was more nuanced. In our clustering analysis, DV cells defined by *sost:NLS-Eos* were primarily observed in cluster 1 and rarely seen in the remaining four clusters. A similar pattern was observed with peripheral cells defined by *sfrp1a:NLS-Eos*, which were usually observed in cluster 0. However, while each marker was specific to certain clusters, these clusters also contained substantial proportions of cells not expressing the marker. This result may indicate that DV and peripheral support cells have a shape phenotype distinct from each other, but only comprise a subset of cells with that shape phenotype. Given that support cells act as multipotent progenitors,^16,32,33^ we speculate that perhaps the more continuous distribution of support cell shapes may reflect this plasticity and smoother transitions between cell states. In contrast, hair cells are a terminally differentiated cell type, which may be represented in their more distinct cell shape phenotype. A caveat is that our cutoff for considering a cell positive for a marker was relatively strict. In addition, our transgenic reporters may not always reflect the cells most recently expressing the transgene due to the time it takes for fluorescent proteins to be expressed and folded and the high stability of the NLS-Eos protein.^34^ Future studies using other methods to label support cell subpopulations with genetic markers, such as RNA-FISH, could further elucidate the connection between gene expression and cell shape in neuromasts. More broadly, cell shape analysis could be integrated with other cellular properties, such as RNA transcript expression, protein expression and organelle location.

We were also interested to see whether nuclear shape could be used to classify cell types. Because the nucleus is often easier to label and segment, nuclear shape representations may be more readily attainable in some developmental biology contexts. We found that nuclear shape could also be used to reliably classify hair cells from support cells. However, compared to cell shape features, nuclear shape features did not appear to distinguish as strongly between the support cell subpopulations we tested in this study. This may suggest that cell shape features are more useful for classifying these populations than nuclear shape features. However, nucleus shape may still be informative in other ways – we observed spatial patterns in nucleus shape within neuromasts, which may represent differences in cell type or state that were not tested in this study. Analysis of nucleus shape may also be more amenable to other tissues where cell membranes may not be easily resolved, allow more rapid collections of larger datasets, or allow analysis of cell types whose cell shapes are not amenable to spherical harmonics analysis. More broadly, we imagine that cell shape, nuclear shape, or a combination of the two may be useful as predictors of cell fate in different tissues.

In many developmental contexts, it remains unclear how genetic factors determine cell shape. We tested whether the gene *atoh1a* regulates cell shape by performing cell shape analysis in *atoh1a* muants. Our results suggest that while *atoh1a* is required for neuromast cells to acquire the hair cell shape phenotype, it may be dispensable for other aspects of neuromast cell shape and organization. We observed that loss of function in *atoh1a* results in the large reduction of one cluster (cluster 4, associated with hair cells) and concurrent increases in existing clusters, but does not lead to generation of a new shape cluster. Therefore, it appears that *atoh1a* is required for the transition from a support cell shape to a hair cell shape; when *atoh1a* function is lost, cells become arrested in shapes that exist in wildtype neuromasts rather than acquiring an *atoh1a* mutant-specific shape phenotype. Similarly, during an RNAi screen for morphological complexity in *Drosophila* Kc cells, RNAi knockdown typically did not generate new shapes but instead changed distributions of pre-existing, wildtype shape states.^35^ It would be illuminating to explore other genes and signaling pathways that may pattern cell shape within neuromasts. For example, Fgf signaling has been shown to be required for apical constriction and rosette formation in the lateral line primordium.^36^ It is possible that Fgf signaling could regulate aspects of neuromast cell shape independently of *atoh1a.* Applying cell shape analysis to query other genetic and pharmacological perturbations could elucidate how cell shape in neuromasts is regulated by different signaling pathways.

Previous work has proposed that hair cell death might induce cell shape changes and tissue deformations that trigger support cells to proliferate; this idea originates from observations that (1) mechanical strain and cellular spreading induce proliferation in some contexts,^4,5,37^ and (2) support cell shape change during hair cell death is positively correlated with an organism’s hair cell regeneration capability.^38–41^ Mechanical cues and cell density might also act as a brake on proliferation during regeneration, similar to what has been observed during development in the murine utricle.^42^ Quantifying cell shape change during hair cell death or targeted ablation of neuromast cells in the presence of reporters for genes of interest could elucidate the role of cell shape in regulating proliferation and regeneration. Of particular interest is Yap, a transcriptional coactivator that regulates cell proliferation downstream of cell adhesions, cell shape change, and mechanical cues.^43^ Yap is known to play a role in regulating proliferation during development and regeneration of hair cells in the mammalian utricle and organ of Corti.^44–46^ Coupling cell shape analysis with reporters for Yap activity and subcellular localization could shed light on whether cell shape change acts upstream of Yap-induced cell proliferation. Cell shape analysis could also be combined with computational simulations and techniques to measure and perturb forces *in vivo* to relate changes in force with cell shape.

Although relating cell shape change to physical forces was previously possible with simpler shape metrics, we anticipate that robust 3D cell shape representations will better detect subtle shape changes and be relevant to a greater variety of tissues than 2D methods.

Previously, Hartmann and colleagues (2020) analyzed 3D cell morphology in the migrating posterior lateral line primordium. Rather than using a spherical harmonics-based shape representation, this study employed a novel method (ISLA-CBE) based on point cloud morphometry. A strength of ISLA-CBE is that rotational variation can be minimized using a pairwise distance transformation, which is arguably less arbitrary than the method of registration used in our study (which tried to preserve biologically meaningful axes while minimizing rotational variation). In addition, ISLA-CBE is not limited to cells that can be accurately spherically parameterized, as ours was (although it is possible this limitation may be overcome with future implementations of spherical harmonics). However, ISLA-CBE does not easily allow cell shapes to be reconstructed from any point in feature space, which is an advantage of our method. We anticipate that both methods will be useful in the future, as the ideal method of cell shape encoding will likely depend on the context and goals of the study.

Although the method of encoding cell shape was different, we observed some commonalities between the results of our study and that of Hartmann and colleagues (2020). Our observations suggested that hair cells represent a discrete morphological state from support cells, while support cells appeared to vary more continuously in shape. Similarly, Hartmann and colleagues (2020) observed a continuous spectrum of shapes, without distinct clusters, in the lateral line primordium. Together, these results suggest a model whereby initially, cells in the lateral line vary continuously in shape, but as hair cells differentiate, they emerge as a discrete shape. How early these patterns develop and whether they change over time or remain static during neuromast deposition and maturity, could be addressed in future studies.

Although our study looked only at homeostatic, mature neuromasts at one time point, we believe the cell shape representation used here could be valuable for integrating information across multiple spatial and temporal scales. Using time lapse imaging and cell tracking, it would be possible to study dynamic cell shape changes during hair cell development, death, and regeneration. We envision that cell shape states could be ordered along trajectories that represent developmental time, analogous to pseudotime analysis employed in single cell RNA sequencing studies.^47^ In addition, shape analysis can be applied to understand the morphologies of whole tissues and organs.^11^ Combining pseudotime analysis with tissue-scale measurements can elucidate the relationship between individual cell shape change and tissue/organ morphogenesis, as has been demonstrated in a study of notochord development in amphioxus.^47^ Quantifying how these relationships evolve over time as the animal grows and develops could shed light into how individual cell shape changes and movements shape organs, as well as how, conversely, the shape of organs constrain individual cell shape. Comparative cell shape studies could also be done between different sensory epithelia in the same organism or even between different species or ages, which could inform evolutionary relationships.

### Limitations of the Study

In the spherical harmonics-based approach used here, all points on the surface of the cell must be able to be mapped onto a sphere to generate accurate reconstructions. However, some of our cells are sufficiently concave that this is not possible, since a line drawn from the center to the outside of the surface would intersect the surface more than once. As a result, 16% of the cells were excluded from further analysis, and some cell shape states may have been lost. Since this problem likely extends to other cell types beyond neuromasts, there is a need for other 3D cell shape analysis methods that can better represent complex, concave shapes. Development and application of alternative methods for spherical parameterization and spherical harmonics expansion offer one potential solution to this problem. For example, Dalmasso and colleagues (2022) represented cell shapes as signed distances from concentric spheres, which enabled them to represent concave structures of mouse limbs using spherical harmonics. Another method using 4D hyperspherical harmonics enables multiple disjointed objects to be parameterized together, and also enables representation of shapes with gaps or holes.^48^ There are also alternatives for data-driven 3D shape representations that are not based on spherical harmonics. Promising methods in this category include 3D point cloud morphometry-based methods like ISLA-CBE^49^ and deep learning approaches such as autoencoders.^50^ With the rate of exciting developments in the field, we expect that many methods will be available to study cell shape in a variety of contexts in the future.

## Materials and methods

### Fish lines

Experiments were conducted on 5 dpf larval zebrafish (*Danio rerio*). The following zebrafish lines were used for these studies: *Tg(-8.0cldnb:LY-EGFP)^zf^*^106*Tg*^ [RRID:ZDB-ALT-060919-2],^19^ *sost^w^*^215*Tg*^ [RRID: ZDB-ALT-190909-10]^16^ *sfrp1a^w^*^217*Tg*^ [RRID:ZDB-ALT-190909-6],^16^ *atoh1a^w^*^271*Tg*^ (this paper). Animals were used at a stage before sex is determined.

### Generation of *atoh1a^w^*^271*Tg*^

The *atoh1a^w^*^271^*^Tg^* line was generated by CRISPR-Cas9 knock-in of the mRuby fluorophore into the coding sequence of the endogenous *atoh1a* locus, as previously published.^16,51^ Homozygotes exhibit phenotypes consistent with *atoh1a* loss of function, such as hearing and vestibular defects, and are viable until approximately 10 days post fertilization (dpf).

### Fish handling

Prior to experiments, embryos and larvae were raised in E3 embryo medium (14.97 mM NaCl, 500 μM KCL, 42 μM NaHPO_4_, 150 μM KH_2_PO_4_, 1 mM CaCl_2_dehydrate, 1 mM MgSO^4^, 0.714 mM NaHCO_3_, pH 7.2) at 28.5°C. Zebrafish larvae were fed rotifers daily from 4 dpf onwards. Experiments used zebrafish at 5 dpf, a stage before sex is determined. Zebrafish experiments and husbandry followed standard protocols in accordance with University of Washington Institutional Animal Care and Use Committee guidelines.

### DRAQ5 staining

On the day of imaging, DRAQ5 dye (Thermo-Fisher) was diluted in E3 embryo medium to a working concentration of 5 µM. 5 dpf larval zebrafish were incubated in this solution for approximately 1 hour, then prepared as described in “Preparation of larvae for live imaging.” We observed best results when the fish were incubated in DRAQ5 no longer than 1 hour and immediately prepared for imaging afterward.

### Photoconversion of NLS-Eos

For experiments done with *sost:NLS-Eos* or *sfrp1a:NLS-Eos* lines, larvae were photoconverted using an an iLumen 8 UV flashlight (purchased from Amazon) for 15-20 minutes. Photoconversion was done by placing fish in a 60 × 15 mm petri dish, removing the petri dish lid, and placing the UV flashlight directly over the dish within a box lined with aluminum foil.

### Preparation of larvae for live imaging

Fish were transferred to E3 embryo medium containing approximately 1.5 mM MESAB to be anesthetized. Once larvae were suitably anesthetized, they were mounted on slides with bridged coverslips in 1.2% low melting point agarose. Fish were mounted in a standard orientation of anterior facing left and dorsal up.

### Confocal imaging

Imaging was performed using a Zeiss LSM 880 with Airyscan equipped with a Zeiss C-Apochromat 40x/1.2W numerical water objective. Imaging was performed at room temperature (approximately 25°C). For each fish, several anterior neuromasts were imaged. Neuromasts were chosen from a list based on which had the most ideal orientations in the fish being imaged (e.g. those neuromasts closest to the coverslip with apical-basal axes roughly parallel to the axial plane/Z-axis of imaging). 633 nm, 561 nm, and 488 nm lasers were used for excitation. Gain was adjusted for different transgenes but kept constant across experiments with the same transgene. Images were collected with physical voxel size of 0.05 x 0.05 x 0.22 µm^3^. All images were captured using the Zen Black acquisition software (Zeiss).

### Image preprocessing

Using Zen Blue, Airyscan Processing was applied to all raw images using the default settings for pixel reassignment and deconvolution.^21^ Next, a custom Python script using the aicsimageio package^52^ was used to remove any unneeded channels and export the CZI files to TIFFs. The original CZI files were kept to ensure access to the original acquisition metadata. To correct for xy drift between z-slices, StackReg was used, either through the MultiStackReg FIJI plugin or a custom script using the pystackreg Python port of StackReg.^53^ Using the “rigid body” method and “previous” setting for the reference, transformation matrices were calculated for the membrane channel, saved, and applied to all channels in the image. Images were manually inspected following stack registration to ensure appropriate drift correction.

Occasionally, imaging acquisition errors or z drift caused either duplication of z slices within the stack or the need to acquire additional z-slices to cover the entire neuromast. In cases where this could be easily manually corrected, duplicated z-slices were deleted, or two z-stacks were concatenated together to cover the entire neuromast. These manual corrections were done using FIJI^54^ prior to stack registration.

### Training of machine learning models for detection of nuclei and membranes

Models to detect neuromast nuclei and membranes were trained using the iterative deep learning workflow of the Allen Cell and Structure Segmenter.^22^ For the membrane model, preliminary segmentations of cell boundaries were initially generated using the lamin B1 workflow from the Segmenter. The “Curator” tool from the Segmenter was used to mask out areas of the images that were not segmented accurately. The original membrane model was trained on a dataset of 14 neuromasts labeled with *Tg(-8.0cldnb:LY-EGFP)*. The membrane model predictions were thresholded and used as input for an interactive distance transform watershed segmentation workflow. Upon generating the label images, the find_boundaries function from scikit-image^57^ was used to generate watershed lines, which was used as input to train a second membrane model with improved performance. This process was repeated once more with a new dataset of 9 *Tg(-8.0cldnb:LY-EGFP)* neuromasts stained with DRAQ5, except labeled nuclei were used to generate seeds for the watershed instead of the distance transform. The results were used to train the third membrane model, which was used for instance segmentation (delineation of individual objects) of all the images in this study.

To generate the nucleus mask model, the “H2B coarse” model provided by AICS^22^ was used to generate preliminary nucleus predictions. The predictions were thresholded, and the membrane intensity image (or membrane boundary predictions) were used to “split” the nuclei (as described in “Instance segmentation”). Similar to the refinement for the membrane model, a distance transform watershed workflow was applied to thresholded nuclei predictions to generate labeled nuclei, which were converted to binary masks and used to train the first nucleus mask model. This process was repeated (starting from the model 1 predictions instead of the H2B coarse model predictions) to generate a second nucleus mask model with improved performance, which was used for all the images in this study.

### Instance segmentation

Instance segmentation of nuclei was done in a semi-automated, interactive fashion using a custom Python script. The Python packages napari^55^ and magicgui^56^ were used to create a graphical user interface for this step, enabling parameter tuning, interactivity, and manual annotations when needed.^55^

First, the nucleus mask predictions and cell boundary predictions were binarized using thresholds of 0.6 and 0.4, respectively. These thresholds were selected a balance between capturing intensity signal corresponding to the structures of interest and minimizing artifacts due to noise. The binarized membrane predictions were used to “split” falsely merged nuclei in the binarized nucleus mask predictions, using one of two methods. For experiments not involving NLS-Eos transgenes, the nucleus mask splitting was done fully automatically (e.g. wherever the membrane mask value was 1, the corresponding pixels in the nucleus mask were set to 0). For experiments involving NLS-Eos transgenes, incomplete photoconversion and/or ongoing transgene expression can cause some signal in the nuclei in the green channel, where the membranes are also imaged. This overlap can create artifacts when the cell boundary model is applied. To circumvent the issue, an alternative “interactive” nucleus splitting strategy was devised. For interactive nucleus splitting, regions of interest (ROIs) were manually drawn slice-by-slice in areas where splitting was desired using the shapes tool in napari. Then, only these regions within the membrane mask were considered during the splitting process.

Following nucleus splitting, individual nuclei were labeled using distance transform watershed. First, the Euclidean distance transform of the nucleus mask was calculated and smoothed using a Gaussian kernel. Next, the local maxima of the smoothed distance transform were calculated using the peak_local_max function from scikit-image.^57^ The local maxima were then used as markers for a watershed segmentation using the inverted distance transform as the input. The sigma parameter for the Gaussian smoothing of the distance transform and the minimum distance parameter for the peak_local_max function were adjusted interactively to yield the best results for each image. Remaining segmentation errors were corrected by manual annotation using a Wacom Cintiq 16 drawing tablet and built-in labels layer tools in napari.^55^

To segment cells, the segmented nuclei were used as markers for a watershed segmentation with the cell boundary predictions (i.e. the raw, unthresholded output of the membrane model inference) as input. The resulting preliminary cell labels were post-processed to remove small objects and manually corrected using a Wacom Cintiq 16 drawing tablet and built-in labels layer tools in napari.^55^

### Cell dataset preparation

Prior to running the cell variance analysis pipeline,^10^ individual cells from the raw and labeled whole neuromast images were cropped, subsampled in z to obtain isotropic voxel size, and saved as individual images. A custom Python script was used to prepare cropped single cells and a manifest.csv file compatible with the pipeline.

### Cell and nucleus alignment

Cells were aligned using a series of three rotations. The first rotation was done with reference to the neuromast organ. The neuromast centroid was calculated from a mask representing the whole neuromast (created from the cell segmentation channel). For each cell, the relative position of the cell centroid to the neuromast centroid was used to calculate the angle between the cell centroid and the x-axis. The corresponding cropped image of the cell was then rotated around the z-axis by this angle to minimize variation originating from the rosette structure of the neuromast.

The second and third rotations were done with reference to the cropped cell only. For the second rotation, the cropped image of the cell was rotated such that the long axis of the cell projected in the xz-plane was aligned to the z-axis. For the second rotation, the cropped image of the cell was rotated such that the long axis of the cell projected in the yz-plane was aligned to the z-axis, unless this angle would be greater than 45 degrees. In that case, the yz long axis was aligned to the y-axis instead. This adjustment prevented cells from being incorrectly rotated by 90 degrees if their long axis was in the horizontal plane.

To generate the nucleus shape space, the nuclei were aligned separately from the cells. The procedure was the same as for the cells, except for the second rotation which also was constrained to be no greater than 45 degrees. If the angle would be greater than 45 degrees, the long axis of the nuclei in the xz-plane was aligned to the x-axis instead of the z-axis.

### Spherical harmonics parameterization

Following cell dataset preparation and alignment, spherical harmonics coefficients for each cell and nucleus were calculated using the cvapipe_analysis package.^10^ First, the “loaddata” step was run with the prepared manifest.csv, then the “computefeatures” step was run. To determine the degree of expansion (L_max_) for spherical harmonics parameterization, we plotted several metrics of reconstruction fidelity for different values of L_max_ using a subset of the data. As L_max_ increases, shapes can be reconstructed with greater detail, but higher L_max_ values incur additional computational costs and exhibit diminishing returns in reconstruction fidelity. We selected an L_max_ of 32 as a value around which reconstruction fidelity plateaued, which yielded 2178 coefficients. The directed Hausdorff distance, which represents the furthest distance between the closest points on the original and reconstructed meshes, was used as a measure of reconstruction error. A Gaussian mixture model was used to identify cells belonging to the second peak in the reconstruction error distribution, which were subsequently excluded from cell shape analysis.

### Principal components analysis and unsupervised clustering

Principal components analysis (PCA) was performed using the Python library scikit-learn.^58^ To create “shape modes,” PCs were z-scored (i.e. the values for each PC were divided by their respective standard deviation). Visualizations of cells in shape space were generated by applying the inverse PCA transform to generate corresponding SHE coefficients, which were then used to generate 3D reconstructions using the aics-shparam package.^10^ Reconstructions were visualized in 3D using ParaView.^59^

Clustering was performed using the Python package Phenograph^25^ with the Leiden algorithm for community detection.^26^ For analysis of cell shape, the first eight PCs were used for clustering; for nuclear shape, the first four PCs were used. These PCs explained 70% and 76% of total variance, respectively. The Uniform Manifold Approximation and Projection (UMAP) implementation from the Python package umap-learn was used to project the data in two dimensions for visualization.^27^ Partition based graph abstraction (PAGA) was performed using the scanpy Python package.^28,60^ PAGA was used to initialize the layout of UMAPs.

### Quantification of fluorescence intensity of transgenes

For reporter lines expressing nuclear localized Eos, *sost:NLS-Eos* and *sfrp1a:NLS-Eos*, the mean intensity within each cell was calculated using the segmented nuclei as masks. For *atoh1a:mRuby*, which has cytoplasmic mRuby localization, the mean intensity within each cell was calculated using the segmented cell as masks. Because intensities of fluorescent proteins from targeted insertions can vary between individual neuromasts and fish, we z-scored the mean intensities for each neuromast and used a threshold of 1 to determine cells “positive” for a marker. In other words, the mean intensity of fluorescent protein within a cell needed to be at least one standard deviation above the mean for that neuromast to be classified as “positive” for that protein.

### Measurement of cell location and shape features

Locations of neuromast cells were estimated using the x and y coordinates of the cell segmentation centroids. The neuromast centroid was calculated by using the cell segmentation channel to generate a mask representing the whole neuromast. For each cell, the distance between the cell centroid and neuromast centroid in the xy plane was calculated. This distance was normalized to that of the cell with the greatest distance from the neuromast centroid to account for differences in neuromast size.^33^

We used the first alignment rotation angle to approximate the angular location of each neuromast cell. This angle represents the angle between the neuromast x-axis (which extends from the neuromast centroid) and a vector drawn from the cell centroid to the neuromast centroid in the xy-plane (Figure 2B). In our images, the neuromast x-axis is approximately parallel to the anterior-posterior body axis of the fish. Lateral line neuromasts can be polarized in one of two ways relative to the anteroposterior body axis: parallel or perpendicular.^61^ Because our dataset contained neuromasts of both types, we standardized the orientations of the neuromasts by adjusting the angles of perpendicular neuromasts using the following formula:

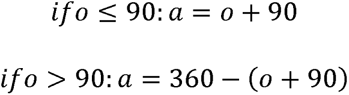

where *o* is the original angle, and *a* is the adjusted angle. This is equivalent to rotating a perpendicular neuromast 90 degrees counterclockwise such that its dorsoventral axis is aligned with the anteroposterior body axis. The decision to rotate counterclockwise was based on the fact that Emx2 is expressed in the anterior region of parallel neuromasts and the dorsal region of perpendicular neuromasts.^62^ The radial distances and angular locations of each cell were combined to plot the locations of neuromast cells on polar coordinates.

Other shape measurements, such as volume, height, etc. were obtained as outputs of the cvapipe_analysis computefeatures step.^10^

### Logistic regression classification

Logistic regression was performed using the Python package scikit-learn.^58^ The data were split into an 80/20 ratio of training data to test data. The test and train subsets were split such that both had similar proportions of hair cells. The logistic regression model was trained to classify hair cells on the train subset, using one or more shape modes as predictor variables. The trained model was then applied to the testing subset to generate the ROC curves and confusion matrices.

### Statistical Analysis

Chi-square analysis was used to distinguish distribution differences among groups, using the Python package SciPy.^63^

## Acknowledgments

The authors thank David White and the UW Fish Facility staff for animal care and maintenance. We thank Julie Theriot and members of her lab for insightful discussion and feedback throughout the course of this study. We also thank Celeste Berg and Cecilia Moens for critical reading of the manuscript.

## Data and Code Availability

Image data is deposited and publicly available at the date of publication at the BioImage Archive (https://www.ebi.ac.uk/bioimage-archive/). All original code is publicly available at the date of publication at https://github.com/raible-lab/neuromast3d.

---

**Supplemental Figure 1.**
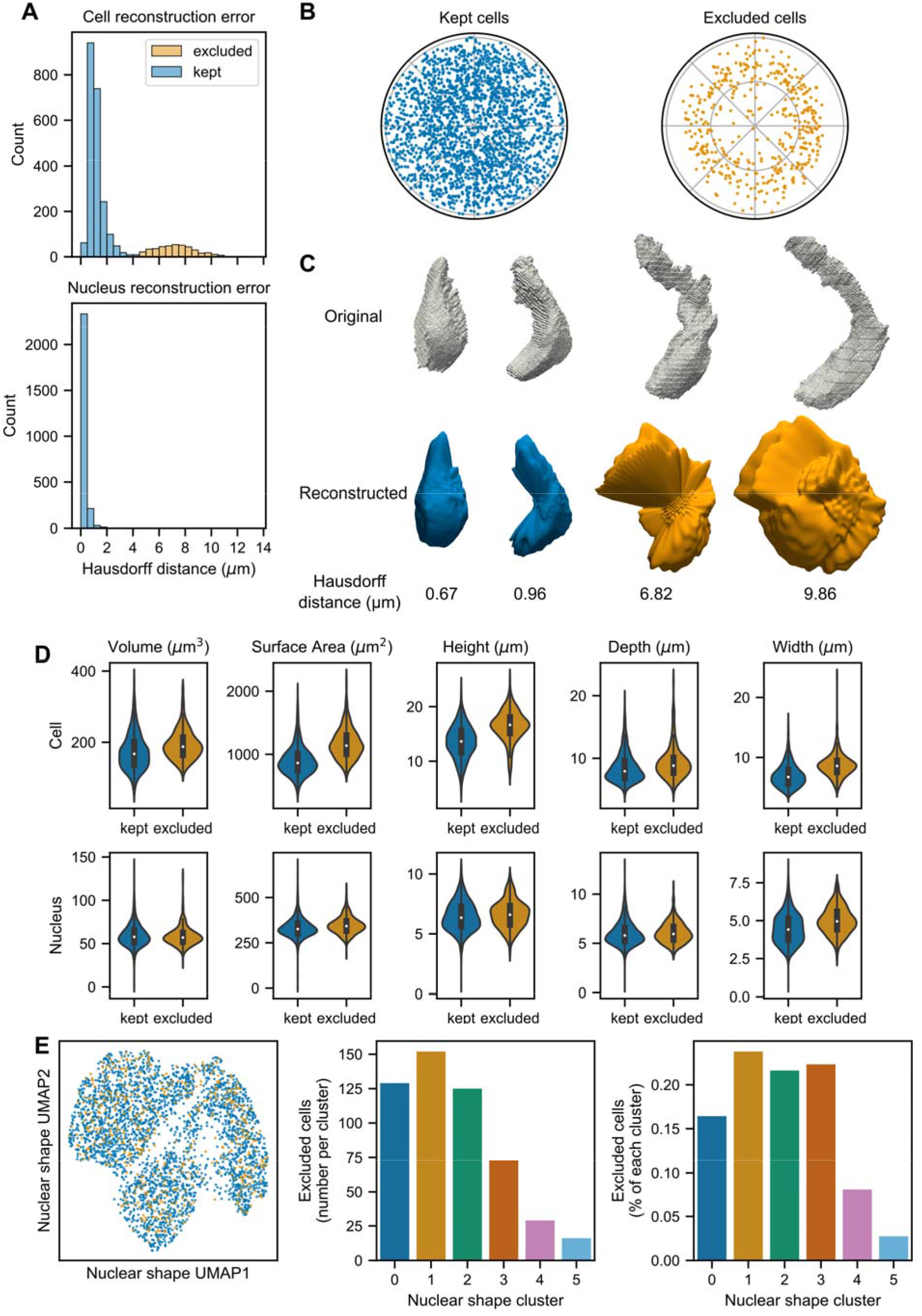
(A) Distribution of reconstruction error, as measured by directed Hausdorff distance, for cell shape (top) and nuclear shape (bottom). Some cells were excluded from cell shape analysis (orange) due to high reconstruction error. (B) Polar plots showing relative locations of cells kept (left, blue) and excluded from cell shape analysis (right, orange). (C) Examples of well reconstructed (left) and poorly reconstructed (right) cells. 3D representations of the original mesh (top), reconstructed mesh (middle), and the associated Hausdorff distances (bottom) for two examples of each are shown. (D) Distributions of cell (top) and nuclei (bottom) shape features in kept (blue) and excluded (orange) cells. (E) Left: UMAP projection of cells in nuclear shape space (corresponding with Figure 5A) colored by whether they were kept (blue) or excluded (orange) from cell shape analysis. Middle: Excluded cells plotted as number (raw counts) for each nuclear shape cluster. Right: Excluded cells as a percentage of each nuclear shape cluster.

**Supplemental Figure 2.**
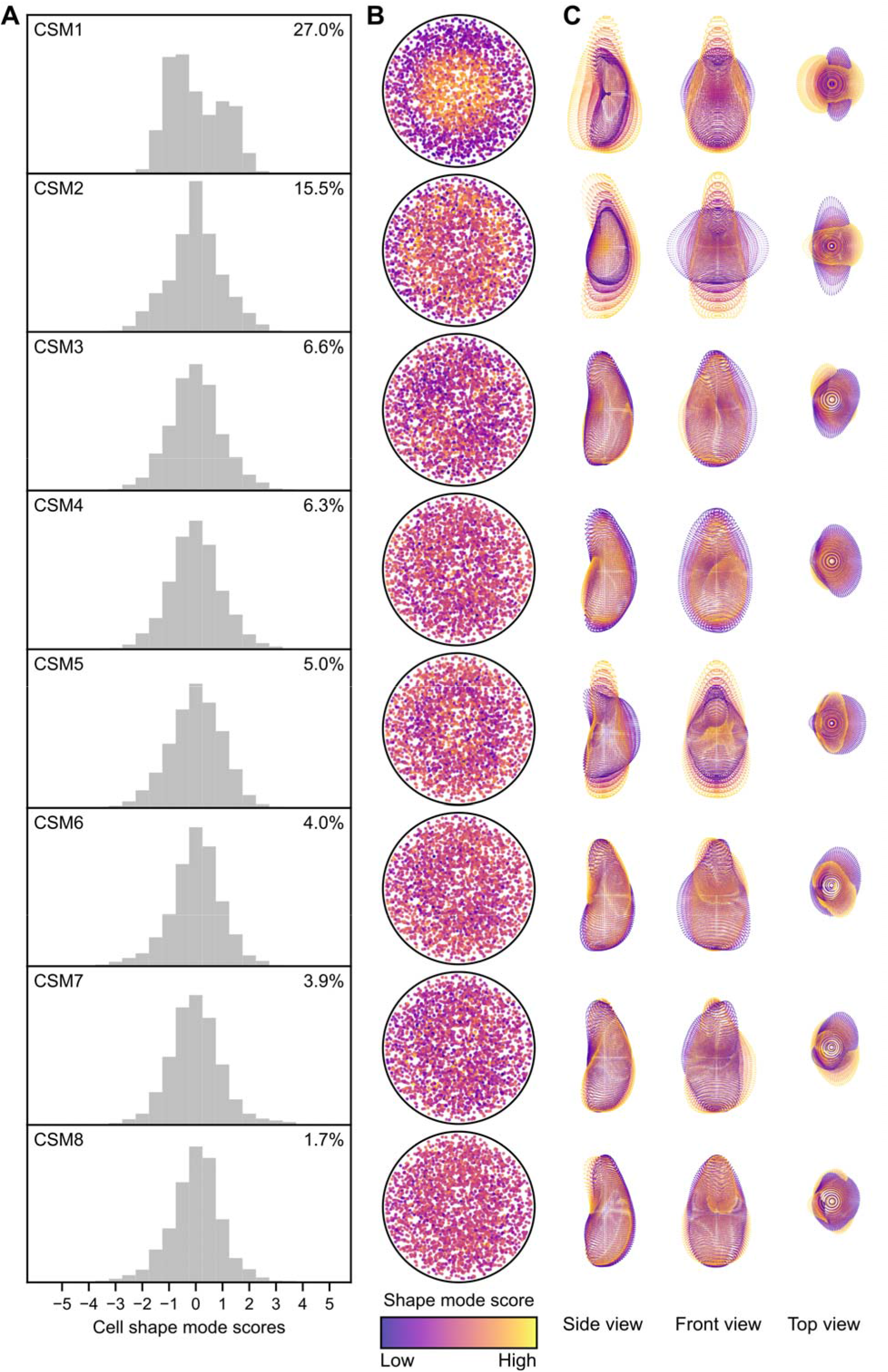
(A) Distributions of shape mode scores for the first eight cell shape modes (CSM1-CSM8). Percent variance explained for each shape mode is provided in the upper right corner. (B) Polar plots showing relative locations of neuromast cells in the dataset, color coded by shape mode scores for CSM1-CSM8. (C) Overlaid 3D point view representations of CSM1-CSM8, generated by finding the mean cell and varying each shape mode by up to 2 standard deviations (sd) in increments of 1 sd while holding other shape modes at the mean (0).

**Supplemental Figure 3.**
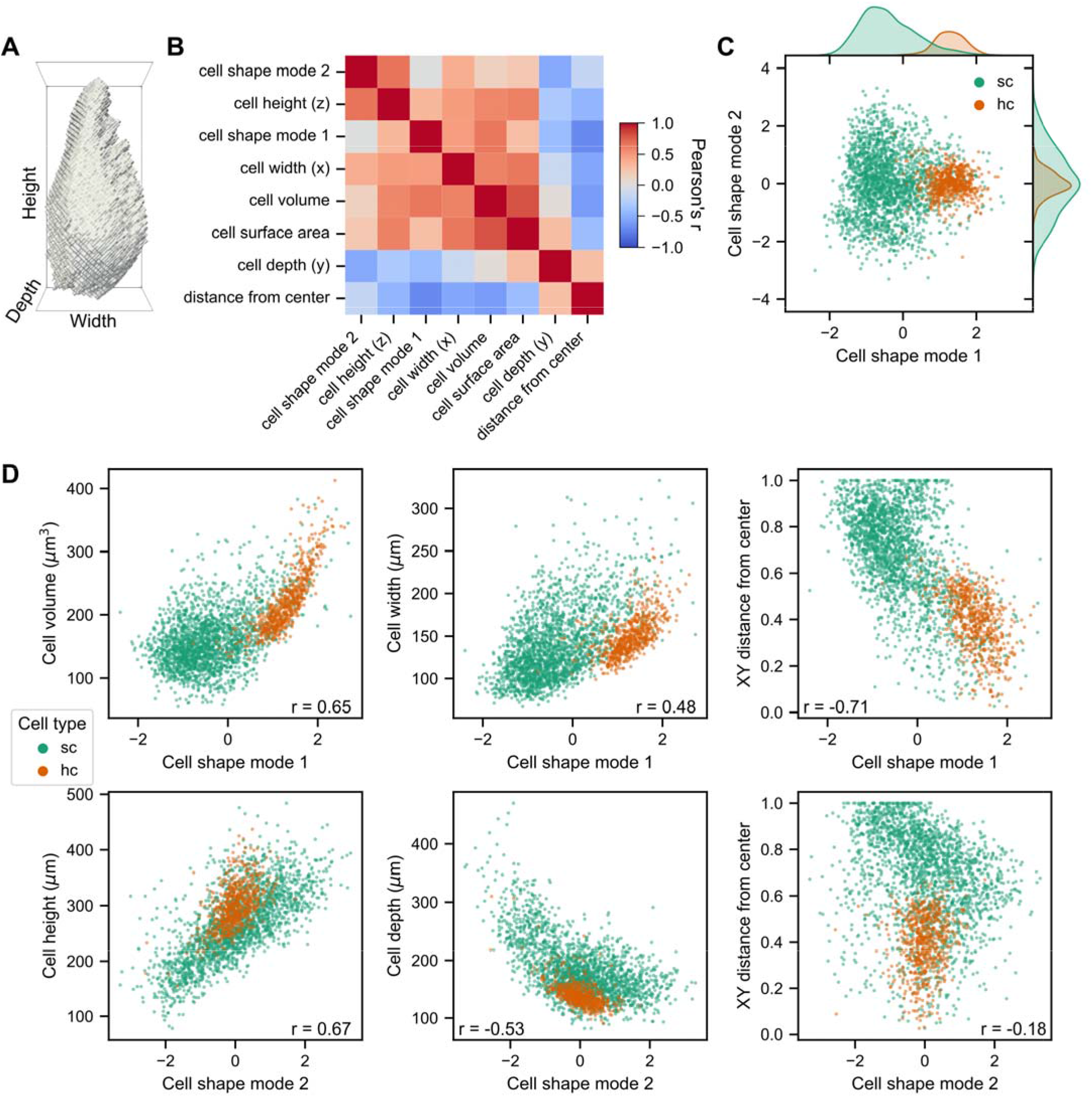
(A) Side view of a 3D projection of a representative cell with its dimensions labeled. (B) Heatmap of correlations between cell shape mode 1, cell shape mode 2, and cell shape/location features. (C) Plot of cell shape mode 1 and cell shape mode 2 for support cells (green) and hair cells (orange). (D) Top row: Correlations between cell volume (left), width (center), and distance from center (right) versus cell shape mode 1 color coded by cell type. Bottom row: Correlations between cell height (left), depth (center), and distance from center (right) versus cell shape mode 2 color coded by cell type. The Pearson correlation coefficient (r) is displayed on each plot. Support cells (green) and hair cells (orange).

**Supplemental Figure 4.**
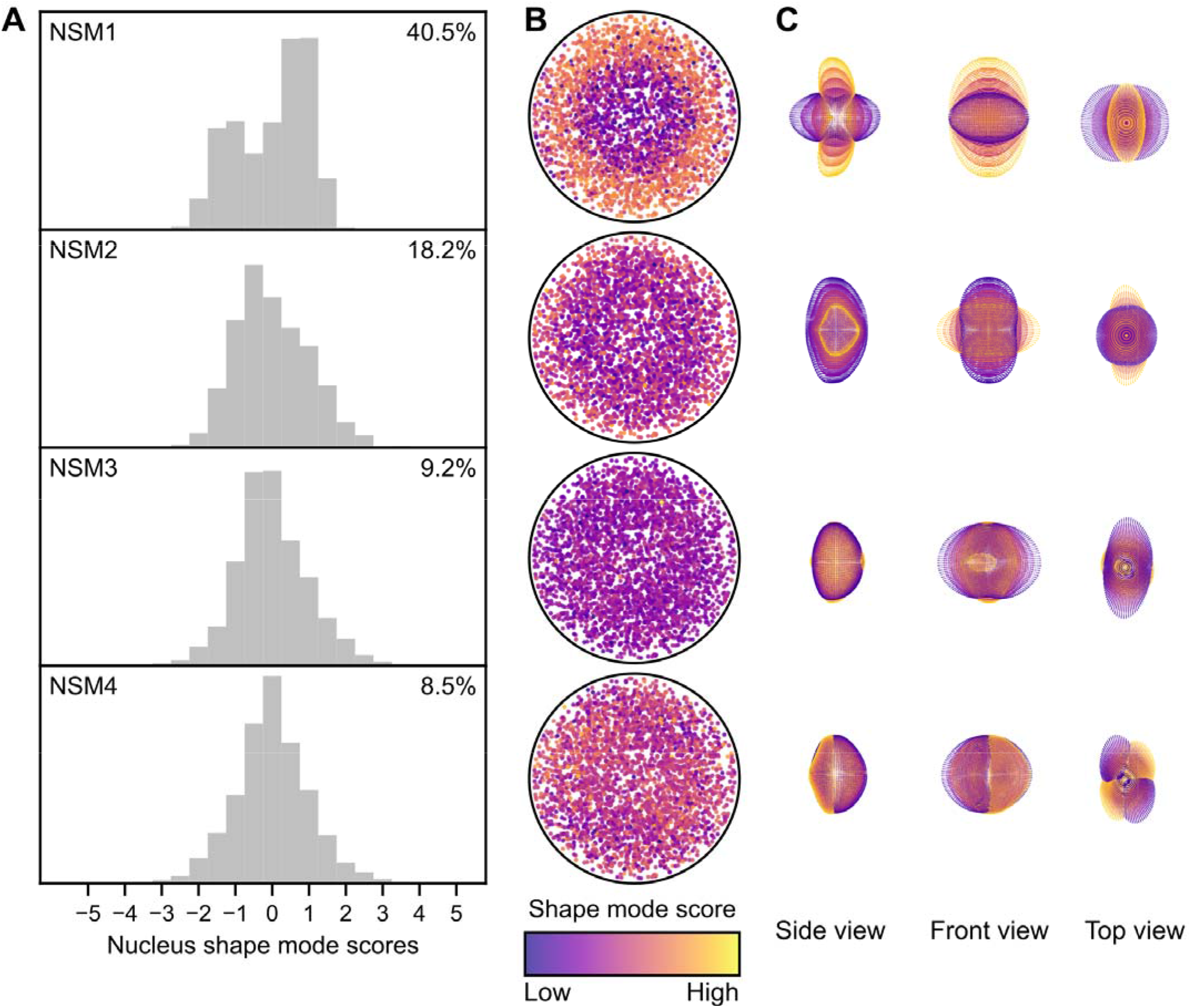
(A) Distributions of shape mode scores for the first four nucleus shape modes (NSM1-NSM4). Percent variance explained for each shape mode is provided in the upper right corner. (B) Polar plots showing relative locations of neuromast cells in the dataset, color coded by shape mode scores for NSM1-NSM4. (C) Overlaid 3D point view representations of CSM1-CSM8, generated by finding the mean cell and varying each shape mode by up to 2 standard deviations (sd) in increments of 1 sd while holding other shape modes at the mean (0).

**Supplemental Figure 5.**
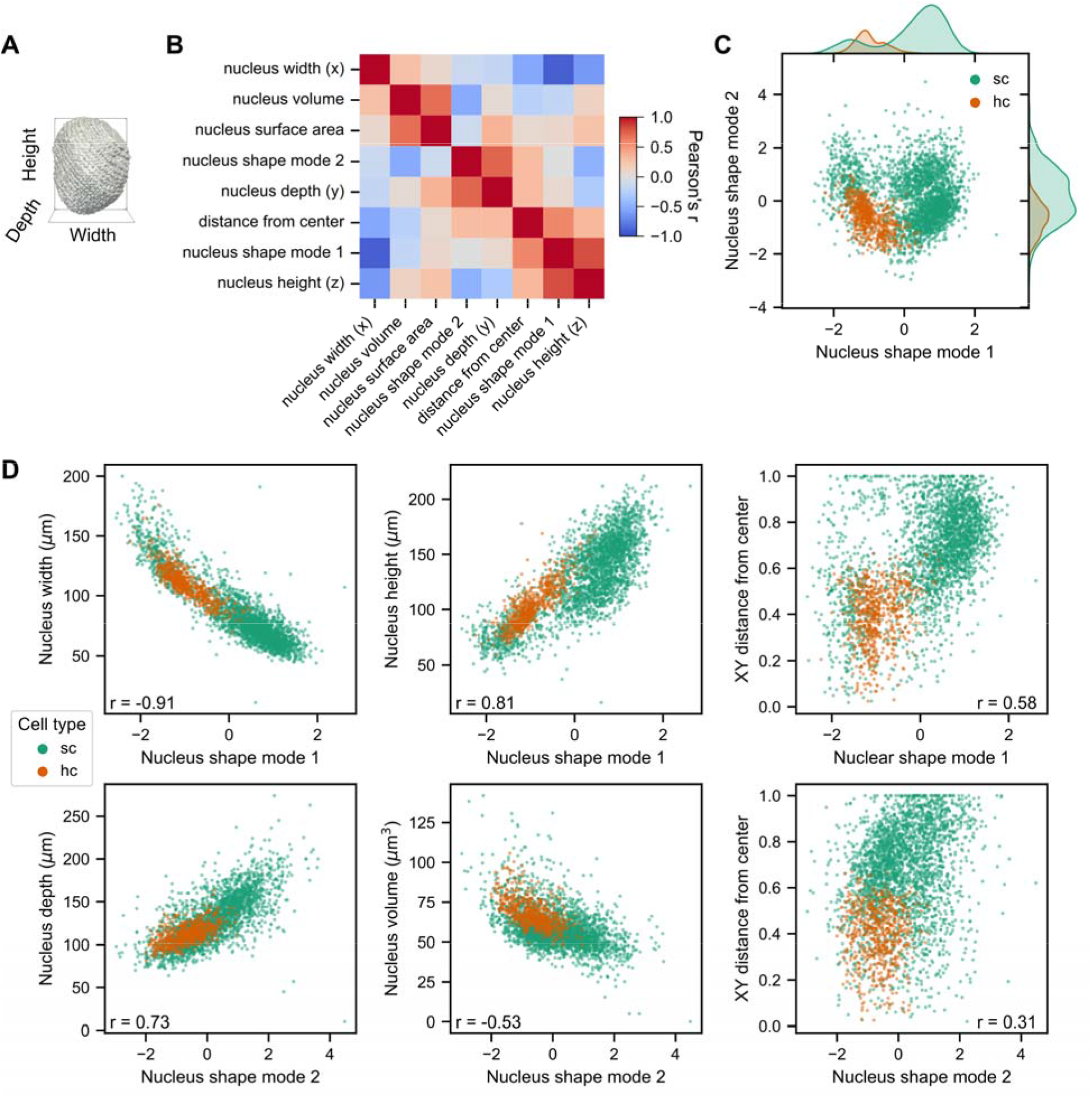
(A) Side view of a 3D projection of a representative nucleus with its dimensions labeled. (B) Heatmap of correlations between nucleus shape mode 1, nucleus shape mode 2, and nucleus shape/location features. (C) Plot of nucleus shape mode 1 and nucleus shape mode 2 for support cells (green) and hair cells (orange). (D) Top row: Correlations between nucleus width (left), height (center), and distance from center (right) versus nucleus shape mode 1 color coded by cell type. Bottom row: Correlations between nucleus depth (left), volume (center), and distance from center (right) versus nucleus shape mode 2 color coded by cell type. The Pearson correlation coefficient (r) is displayed on each plot. Support cells (green) and hair cells (orange).

**Supplemental Figure 6.**
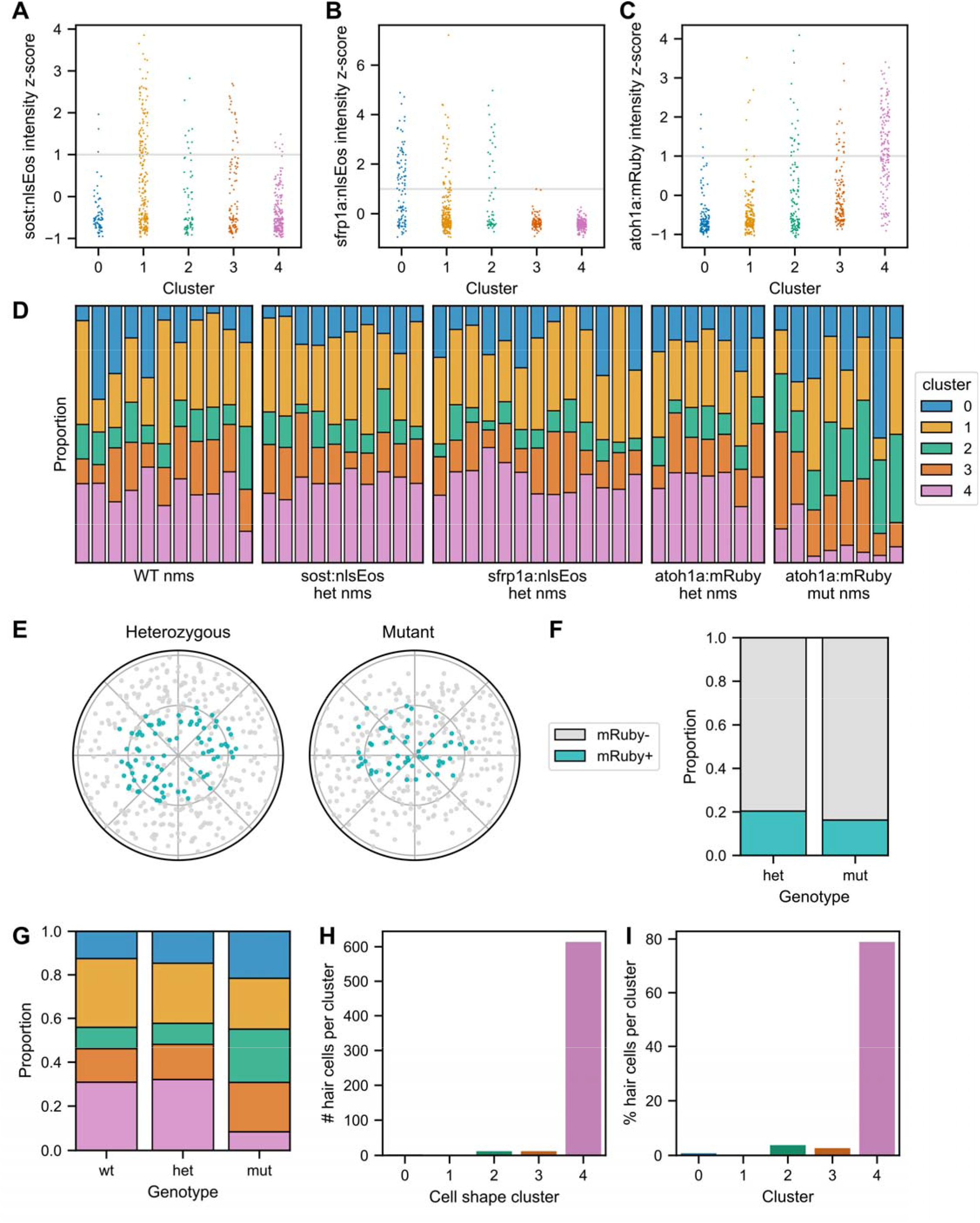
(A) Distribution of mean *sost:NLS-Eos* intensities (z-scored) for cells in each cell shape cluster. Gray line represents intensity threshold (1 SD) used to label a cell as positive. (B) Distribution of mean *sfrp1a:NLS-Eos* intensities (z-scored) for cells in each cell shape cluster. Gray line represents intensity threshold (1 SD) used to label a cell as positive. (C) Distribution of mean *atoh1a:mRuby* intensities (z-scored) for cells in each cell shape cluster. Gray line represents intensity threshold (1 SD) used to label a cell as positive. (D) Proportions of cells in individual neuromasts belonging to each cluster, grouped by reporter expression and genotype (n=49 neuromasts). (E) Polar plots of relative cell locations of mRuby-cells (gray) and mRuby+ cells (dark cyan) in *atoh1a:mRuby* heterozygotes (left) and *atoh1a:mRuby* mutants (right). (F) Proportions of mRuby+ cells (dark cyan) and mRuby-cells (gray) in *atoh1a:mRuby* heterozygotes (left bar) and *atoh1a:mRuby* mutants (right bar). (G) Proportions of cells of each genotype - *atoh1a:mRuby* mutants, heterozygotes, and wildtype - belonging to each cell shape cluster. (H) Hair cells as raw counts (number) within each cell shape cluster. (I) Hair cells as a percentage of each cell shape cluster.

**Supplemental Table 1.** Summary of the dataset. Each batch represents a separate imaging session. All neuromasts were labeled with DRAQ5 dye in the *Tg(-8.0cldnb:LY-EGFP)* background; the “reporter” column indicates additional genetic labels. “Filter” refers to cells excluded for having high cell shape reconstruction error (see Supplemental Figure 1 for details).

| batch | reporter | # neuromasts | # cells (pre-filter) | # cells (post-filter) |
| --- | --- | --- | --- | --- |
| 1 | sfrp1a:nlsEos | 7 | 453 | 383 |
| 2 | sost:nlsEos | 10 | 824 | 676 |
| 3 | sfrp1a:nlsEos | 6 | 459 | 371 |
| 4 | atoh1a:mRuby | 10 | 560 | 496 |
| 5 | atoh1a:mRuby | 5 | 295 | 244 |
| 6 | N/A (wildtype) | 11 | 683 | 580 |

## References

1. Chan, C.J., Heisenberg, C.-P., and Hiiragi, T. (2017). Coordination of Morphogenesis and Cell-Fate Specification in Development. Curr. Biol. 27, R1024–R1035. 10.1016/j.cub.2017.07.010.

2. Butler, L.C., Blanchard, G.B., Kabla, A.J., Lawrence, N.J., Welchman, D.P., Mahadevan, L., Adams, R.J., and Sanson, B. (2009). Cell shape changes indicate a role for extrinsic tensile forces in Drosophila germ-band extension. Nat. Cell Biol. 11, 859–864. 10.1038/ncb1894.

3. Sun, Z., Amourda, C., Shagirov, M., Hara, Y., Saunders, T.E., and Toyama, Y. (2017). Basolateral protrusion and apical contraction cooperatively drive Drosophila germ-band extension. Nat. Cell Biol. 19, 375–383. 10.1038/ncb3497.

4. Chen, C.S., Mrksich, M., Huang, S., Whitesides, G.M., and Ingber, D.E. (1997). Geometric Control of Cell Life and Death. Science 276, 1425–1428. 10.1126/science.276.5317.1425.

5. Watt, F.M., Jordan, P.W., and O’Neill, C.H. (1988). Cell shape controls terminal differentiation of human epidermal keratinocytes. Proc. Natl. Acad. Sci. 85, 5576–5580. 10.1073/pnas.85.15.5576.

6. Kilian, K.A., Bugarija, B., Lahn, B.T., and Mrksich, M. (2010). Geometric cues for directing the differentiation of mesenchymal stem cells. Proc. Natl. Acad. Sci. 107, 4872–4877. 10.1073/pnas.0903269107.

7. Pincus, Z., and Theriot, J.A. (2007). Comparison of quantitative methods for cell-shape analysis. J. Microsc. 227, 140–156. 10.1111/j.1365-2818.2007.01799.x.

8. Keren, K., Pincus, Z., Allen, G.M., Barnhart, E.L., Marriott, G., Mogilner, A., and Theriot, J.A. (2008). Mechanism of shape determination in motile cells. Nature 453, 475–480. 10.1038/nature06952.

9. Ruan, X., and Murphy, R.F. (2019). Evaluation of methods for generative modeling of cell and nuclear shape. Bioinformatics 35, 2475–2485. 10.1093/bioinformatics/bty983.

10. Viana, M.P., Chen, J., Knijnenburg, T.A., Vasan, R., Yan, C., Arakaki, J.E., Bailey, M., Berry, B., Borensztejn, A., Brown, E.M., et al. (2023). Integrated intracellular organization and its variations in human iPS cells. Nature 613, 345–354. 10.1038/s41586-022-05563-7.

11. Dalmasso, G., Musy, M., Niksic, M., Robert-Moreno, A., Badía-Careaga, C., Sanz-Ezquerro, J.J., and Sharpe, J. (2022). 4D reconstruction of murine developmental trajectories using spherical harmonics. Dev. Cell 57, 2140–2150.e5. 10.1016/j.devcel.2022.08.005.

12. Gerig, G., Styner, M., Jones, D., Weinberger, D., and Lieberman, J. (2001). Shape analysis of brain ventricles using SPHARM. In Proceedings IEEE Workshop on Mathematical Methods in Biomedical Image Analysis (MMBIA 2001), pp. 171–178. 10.1109/MMBIA.2001.991731.

13. López-Schier, H., and Hudspeth, A.J. (2006). A two-step mechanism underlies the planar polarization of regenerating sensory hair cells. Proc. Natl. Acad. Sci. 103, 18615–18620. 10.1073/pnas.0608536103.

14. Pinto-Teixeira, F., Viader-Llargués, O., Torres-Mejía, E., Turan, M., González-Gualda, E., Pola-Morell, L., and López-Schier, H. (2015). Inexhaustible hair-cell regeneration in young and aged zebrafish. Biol. Open 4, 903–909. 10.1242/bio.012112.

15. Lush, M.E., Diaz, D.C., Koenecke, N., Baek, S., Boldt, H., St Peter, M.K., Gaitan-Escudero, T., Romero-Carvajal, A., Busch-Nentwich, E.M., Perera, A.G., et al. (2019). scRNA-Seq reveals distinct stem cell populations that drive hair cell regeneration after loss of Fgf and Notch signaling. eLife 8, e44431. 10.7554/eLife.44431.

16. Thomas, E.D., and Raible, D.W. (2019). Distinct progenitor populations mediate regeneration in the zebrafish lateral line. eLife 8, e43736. 10.7554/eLife.43736.

17. Bermingham, N.A., Hassan, B.A., Price, S.D., Vollrath, M.A., Ben-Arie, N., Eatock, R.A., Bellen, H.J., Lysakowski, A., and Zoghbi, H.Y. (1999). Math1: An Essential Gene for the Generation of Inner Ear Hair Cells. Science 284, 1837–1841. 10.1126/science.284.5421.1837.

18. Millimaki, B.B., Sweet, E.M., Dhason, M.S., and Riley, B.B. (2007). Zebrafish atoh1 genes: classic proneural activity in the inner ear and regulation by Fgf and Notch. Development 134, 295–305. 10.1242/dev.02734.

19. Haas, P., and Gilmour, D. (2006). Chemokine Signaling Mediates Self-Organizing Tissue Migration in the Zebrafish Lateral Line. Dev. Cell 10, 673–680. 10.1016/j.devcel.2006.02.019.

20. Smith, P.J., Wiltshire, M., Davies, S., Patterson, L.H., and Hoy, T. (1999). A novel cell permeant and far red-fluorescing DNA probe, DRAQ5, for blood cell discrimination by flow cytometry. J. Immunol. Methods 229, 131–139. 10.1016/s0022-1759(99)00116-7.

21. Huff, J. (2015). The Airyscan detector from ZEISS: confocal imaging with improved signal-to-noise ratio and super-resolution. Nat. Methods 12, i–ii. 10.1038/nmeth.f.388.

22. Chen, J., Ding, L., Viana, M.P., Lee, H., Sluezwski, M.F., Morris, B., Hendershott, M.C., Yang, R., Mueller, I.A., and Rafelski, S.M. (2018). The Allen Cell and Structure Segmenter: a new open source toolkit for segmenting 3D intracellular structures in fluorescence microscopy images (Cell Biology) 10.1101/491035.

23. Beucher, S., and Lantuéjoul, C. (1979). Use of Watersheds in Contour Detection. In Proceedings of the International workshop on image processing, real-time edge and motion detection (CCETT).

24. Huttenlocher, D.P., Klanderman, G.A., and Rucklidge, W.J. (1993). Comparing images using the Hausdorff distance. IEEE Trans. Pattern Anal. Mach. Intell. 15, 850–863. 10.1109/34.232073.

25. Levine, J.H., Simonds, E.F., Bendall, S.C., Davis, K.L., Amir, E.D., Tadmor, M.D., Litvin, O., Fienberg, H.G., Jager, A., Zunder, E.R., et al. (2015). Data-Driven Phenotypic Dissection of AML Reveals Progenitor-like Cells that Correlate with Prognosis. Cell 162, 184–197. 10.1016/j.cell.2015.05.047.

26. Traag, V.A., Waltman, L., and van Eck, N.J. (2019). From Louvain to Leiden: guaranteeing well-connected communities. Sci. Rep. 9, 5233. 10.1038/s41598-019-41695-z.

27. McInnes, L., Healy, J., Saul, N., and Großberger, L. (2018). UMAP: Uniform Manifold Approximation and Projection. J. Open Source Softw. 3, 861. 10.21105/joss.00861.

28. Wolf, F.A., Hamey, F.K., Plass, M., Solana, J., Dahlin, J.S., Göttgens, B., Rajewsky, N., Simon, L., and Theis, F.J. (2019). PAGA: graph abstraction reconciles clustering with trajectory inference through a topology preserving map of single cells. Genome Biol. 20, 59. 10.1186/s13059-019-1663-x.

29. Itoh, M., and Chitnis, A.B. (2001). Expression of proneural and neurogenic genes in the zebrafish lateral line primordium correlates with selection of hair cell fate in neuromasts. Mech. Dev. 102, 263–266. 10.1016/S0925-4773(01)00308-2.

30. Marklein, R.A., Lo Surdo, J.L., Bellayr, I.H., Godil, S.A., Puri, R.K., and Bauer, S.R. (2016). High Content Imaging of Early Morphological Signatures Predicts Long Term Mineralization Capacity of Human Mesenchymal Stem Cells upon Osteogenic Induction. Stem Cells 34, 935–947. 10.1002/stem.2322.

31. Gerbin, K.A., Grancharova, T., Donovan-Maiye, R.M., Hendershott, M.C., Anderson, H.G., Brown, J.M., Chen, J., Dinh, S.Q., Gehring, J.L., Johnson, G.R., et al. (2021). Cell states beyond transcriptomics: Integrating structural organization and gene expression in hiPSC-derived cardiomyocytes. Cell Syst. 12, 670–687.e10. 10.1016/j.cels.2021.05.001.

32. Romero-Carvajal, A., Navajas Acedo, J., Jiang, L., Kozlovskaja-Gumbrienė, A., Alexander, R., Li, H., and Piotrowski, T. (2015). Regeneration of Sensory Hair Cells Requires Localized Interactions between the Notch and Wnt Pathways. Dev. Cell 34, 267–282. 10.1016/j.devcel.2015.05.025.

33. Viader-Llargués, O., Lupperger, V., Pola-Morell, L., Marr, C., and López-Schier, H. (2018). Live cell-lineage tracing and machine learning reveal patterns of organ regeneration. eLife 7, e30823. 10.7554/eLife.30823.

34. Cruz, I.A., Kappedal, R., Mackenzie, S.M., Hailey, D.W., Hoffman, T.L., Schilling, T.F., and Raible, D.W. (2015). Robust regeneration of adult zebrafish lateral line hair cells reflects continued precursor pool maintenance. Dev. Biol. 402, 229–238. 10.1016/j.ydbio.2015.03.019.

35. Yin, Z., Sadok, A., Sailem, H., McCarthy, A., Xia, X., Li, F., Garcia, M.A., Evans, L., Barr, A.R., Perrimon, N., et al. (2013). A screen for morphological complexity identifies regulators of switch-like transitions between discrete cell shapes. Nat. Cell Biol. 15, 860–871. 10.1038/ncb2764.

36. Harding, M.J., and Nechiporuk, A.V. (2012). Fgfr-Ras-MAPK signaling is required for apical constriction via apical positioning of Rho-associated kinase during mechanosensory organ formation. Development 139, 3130–3135. 10.1242/dev.082271.

37. Benham-Pyle, B.W., Pruitt, B.L., and Nelson, W.J. (2015). Mechanical strain induces E-cadherin–dependent Yap1 and β-catenin activation to drive cell cycle entry. Science 348, 1024–1027. 10.1126/science.aaa4559.

38. Meyers, J.R., and Corwin, J.T. (2007). Shape Change Controls Supporting Cell Proliferation in Lesioned Mammalian Balance Epithelium. J. Neurosci. 27, 4313–4325. 10.1523/JNEUROSCI.5023-06.2007.

39. Burns, J., Christophel, J.J., Collado, M.S., Magnus, C., Carfrae, M., and Corwin, J.T. (2008). Reinforcement of cell junctions correlates with the absence of hair cell regeneration in mammals and its occurrence in birds. J. Comp. Neurol. 511, 396–414. 10.1002/cne.21849.

40. Burns, J.C., Collado, M.S., Oliver, E.R., and Corwin, J.T. (2013). Specializations of intercellular junctions are associated with the presence and absence of hair cell regeneration in ears from six vertebrate classes. J. Comp. Neurol. 521, 1430–1448. 10.1002/cne.23250.

41. Burns, J.C., and Corwin, J.T. (2014). Responses to Cell Loss Become Restricted as the Supporting Cells in Mammalian Vestibular Organs Grow Thick Junctional Actin Bands That Develop High Stability. J. Neurosci. 34, 1998–2011. 10.1523/JNEUROSCI.4355-13.2014.

42. Gnedeva, K., Jacobo, A., Salvi, J.D., Petelski, A.A., and Hudspeth, A.J. (2017). Elastic force restricts growth of the murine utricle. eLife 6, e25681. 10.7554/eLife.25681.

43. Piccolo, S., Dupont, S., and Cordenonsi, M. (2014). The Biology of YAP/TAZ: Hippo Signaling and Beyond. Physiol. Rev. 10.1152/physrev.00005.2014.

44. Rudolf, M.A., Andreeva, A., Kozlowski, M.M., Kim, C.E., Moskowitz, B.A., Anaya-Rocha, A., Kelley, M.W., and Corwin, J.T. (2020). YAP Mediates Hair Cell Regeneration in Balance Organs of Chickens, But LATS Kinases Suppress Its Activity in Mice. J. Neurosci. 40, 3915– 3932. 10.1523/JNEUROSCI.0306-20.2020.

45. Gnedeva, K., Wang, X., McGovern, M.M., Barton, M., Tao, L., Trecek, T., Monroe, T.O., Llamas, J., Makmura, W., Martin, J.F., et al. (2020). Organ of Corti size is governed by Yap/Tead-mediated progenitor self-renewal. Proc. Natl. Acad. Sci. 117, 13552–13561. 10.1073/pnas.2000175117.

46. Kastan, N., Gnedeva, K., Alisch, T., Petelski, A.A., Huggins, D.J., Chiaravalli, J., Aharanov, A., Shakked, A., Tzahor, E., Nagiel, A., et al. (2021). Small-molecule inhibition of Lats kinases may promote Yap-dependent proliferation in postmitotic mammalian tissues. Nat. Commun. 12, 3100. 10.1038/s41467-021-23395-3.

47. Andrews, T.G.R., Pönisch, W., Paluch, E.K., Steventon, B.J., and Benito-Gutierrez, E. (2021). Single-cell morphometrics reveals ancestral principles of notochord development. Development 148, dev199430. 10.1242/dev.199430.

48. Pasha Hosseinbor, A., Chung, M.K., Koay, C.G., Schaefer, S.M., van Reekum, C.M., Schmitz, L.P., Sutterer, M., Alexander, A.L., and Davidson, R.J. (2015). 4D hyperspherical harmonic (HyperSPHARM) representation of surface anatomy: A holistic treatment of multiple disconnected anatomical structures. Med. Image Anal. 22, 89–101. 10.1016/j.media.2015.02.004.

49. Hartmann, J., Wong, M., Gallo, E., and Gilmour, D. (2020). An image-based data-driven analysis of cellular architecture in a developing tissue. eLife 9, e55913. 10.7554/eLife.55913.

50. De Vries, M., Dent, L., Curry, N., Rowe-Brown, L., Bousgouni, V., Tyson, A., Dunsby, C., and Bakal, C. (2022). 3D single-cell shape analysis using geometric deep learning (Bioinformatics) 10.1101/2022.06.17.496550.

51. Kimura, Y., Hisano, Y., Kawahara, A., and Higashijima, S. (2014). Efficient generation of knock-in transgenic zebrafish carrying reporter/driver genes by CRISPR/Cas9-mediated genome engineering. Sci. Rep. 4, srep06545. 10.1038/srep06545.

52. Brown, E.M., Toloudis, D., Sherman, J., Swain-Bowden, M., Lambert, T., and AICSImageIO Contributors (2021). AICSImageIO: Image Reading, Metadata Conversion, and Image Writing for Microscopy Images in Pure Python.

53. Thevenaz, P., Ruttimann, U.E., and Unser, M. (1998). A pyramid approach to subpixel registration based on intensity. IEEE Trans. Image Process. 7, 27–41. 10.1109/83.650848.

54. Schindelin, J., Arganda-Carreras, I., Frise, E., Kaynig, V., Longair, M., Pietzsch, T., Preibisch, S., Rueden, C., Saalfeld, S., Schmid, B., et al. (2012). Fiji: an open-source platform for biological-image analysis. Nat. Methods 9, 676–682. 10.1038/nmeth.2019.

55. van der Walt, S., Schönberger, J.L., Nunez-Iglesias, J., Boulogne, F., Warner, J.D., Yager, N., Gouillart, E., and Yu, T. (2014). scikit-image: image processing in Python. PeerJ 2, e453. 10.7717/peerj.453.

56. Sofroniew, N., Lambert, T., Evans, K., Nunez-Iglesias, J., Bokota, G., Winston, P., Peña-Castellanos, G., Yamauchi, K., Bussonnier, M., Doncila Pop, D., et al. (2022). napari: a multi-dimensional image viewer for Python. 10.5281/ZENODO.7276432.

57. Lambert, T., Liu, H., Buckley, G., Bokota, G., Stansby, D., Har-Gil, H., Nunez-Iglesias, J., Weigert, M., aeisenbarth, Selzer, G., et al. (2023). pyapp-kit/magicgui: v0.7.2. 10.5281/zenodo.7700260.

58. Pedregosa, F., Varoquaux, G., Gramfort, A., Michel, V., Thirion, B., Grisel, O., Blondel, M., Prettenhofer, P., Weiss, R., Dubourg, V., et al. (2011). Scikit-learn: Machine Learning in Python. J. Mach. Learn. Res. 12, 2825–2830.

59. Ahrens, J., Geveci, B., and Law, C. (2005). ParaView: An End-User Tool for Large-Data Visualization. In Visualization Handbook (Elsevier), pp. 717–731. 10.1016/B978-012387582-2/50038-1.

60. Wolf, F.A., Angerer, P., and Theis, F.J. (2018). SCANPY: large-scale single-cell gene expression data analysis. Genome Biol. 19, 15. 10.1186/s13059-017-1382-0.

61. López-Schier, H., Starr, C.J., Kappler, J.A., Kollmar, R., and Hudspeth, A.J. (2004). Directional Cell Migration Establishes the Axes of Planar Polarity in the Posterior Lateral-Line Organ of the Zebrafish. Dev. Cell 7, 401–412. 10.1016/j.devcel.2004.07.018.

62. Jiang, T., Kindt, K., and Wu, D.K. (2017). Transcription factor Emx2 controls stereociliary bundle orientation of sensory hair cells. eLife. 10.7554/eLife.2366

63. Virtanen, P., Gommers, R., Oliphant, T.E., Haberland, M., Reddy, T., Cournapeau, D., Burovski, E., Peterson, P., Weckesser, W., Bright, J., van der Walt, S.J., Brett, M., Wilson, J., Millman, K.J., Mayorov, N., Nelson, A.R.J., Jones, E., Kern, R., Larson, E., Carey, C.J., Polat, İ., Feng, Y., Moore, E.W., VanderPlas, J., Laxalde, D., Perktold, J., Cimrman, R., Henriksen, I., Quintero, E.A., Harris, C.R., Archibald, A.M., Ribeiro, A.H., Pedregosa, F., van Mulbregt, P.; SciPy 1.0 Contributors. (2020) SciPy 1.0: fundamental algorithms for scientific computing in Python. Nat Methods. 17, 261–272. 10.1038/s41592-019-0686-2.

